# High calcium to sheep and goats antepartum Antepartum high dietary supply of calcium affects bone homeostasis and offspring growth in dairy sheep and dairy goats

**DOI:** 10.1101/2024.08.12.607180

**Authors:** D. Brugger, A. Liesegang

## Abstract

**Interpretive Summary:** Many studies have focused on the effects of deficient Ca supply in small dairy ruminants, while little is known about the impact of high dietary Ca intake during the last days before parturition. Here, we show that multiparous dairy goats and sheep maintained Ca homeostasis despite receiving a high dietary Ca supply, with no adverse consequences for their health. However, we observed that animals fed the high dietary Ca diet antepartum showed a gradual increase in serum osteocalcin postpartum, and their suckling offspring exhibited reduced average daily weight gain. The functional background and practical consequences of these findings are yet unknown and may guide future research on the matter.

This study investigated the effects of high dietary Ca supplementation during the final 21 days antepartum on Ca and bone homeostasis in dairy sheep and goats, and the growth response of their suckling offspring. Multiparous dairy sheep (n = 5/group, 10 animals total) and goats (n = 6/group, 12 animals total) were randomly assigned to 2 experimental groups. Feeding occurred restrictively (2.3 kg DM/d*animal^-1^ antepartum for both species; 2.9 and 3.1 kg DM/d*animal^-1^ for sheep and goats postpartum, respectively) according to recommendations, except for Ca during antepartum feeding: 1 group received the basal diet based on hay and concentrate with extra CaCO_3_ (1.3% Ca in DM; 2.49-fold the recommendations) for 3 weeks antepartum, while the other group received just the basal diet (0.6% Ca in DM; 1.15-fold the recommendations). Experimental feeding ended with parturition and animals were henceforth fed according to Swiss recommendations for lactating sheep and goats and kept together with their suckling offspring. The observation period spanned from 21 days antepartum to 56 days postpartum. Animals were under continuous veterinary surveillance and monitored for signs of milk fever. Data collection comprised quantitative and functional parameters of Ca and bone homeostasis as well as birth weights and daily weight gain of suckling lambs and kids. Data were analyzed using repeated measures and endpoint mixed models. The response of quantitative markers indicated that high dietary Ca antepartum significantly increased fecal Ca concentrations until parturition, suggesting efficient physiological mechanisms to manage Ca overload through increasing fecal excretion. No significant differences were observed in serum Ca levels, urinary Ca excretion, or bone mineral density between the antepartum Ca feeding groups at any point during the observation period, indicating stable Ca homeostasis despite the dietary challenge. Differences in quantitative markers were noted between sheep and goats, including variations in serum and colostral Ca levels and bone mineral density, which largely aligned with results from earlier comparative studies. Serum 1,25-(OH)_2_ vitamin D (**calcitriol**) as well as markers of bone formation and resorption were monitored, revealing significant increases in serum osteocalcin postpartum in both goats and sheep fed high Ca antepartum. However, all other serum markers including calcitriol, remained unaffected by the feeding regime but differed between sheep and goats, consistent with previous findings. All repeated measures were significantly affected by time, except for urinary Ca and bone-specific alkaline phosphatase activity in serum. Suckling offspring of sheep and goats in the high Ca group exhibited significantly reduced average daily weight gain compared to those in the 0.6 % in DM Ca group, despite similar birth weights. We conclude that while dairy sheep and goats effectively managed high dietary Ca intake without overt signs of hypocalcemia or milk fever under these experimental conditions, the observed impacts on offspring growth and potential long-term physiological effects warrant further investigation. These findings contribute to the understanding of mineral nutrition in late-gestating dairy goats and sheep and highlight the need for further research on balanced dietary strategies to optimize health and productivity of dairy sheep and goats and their offspring. Key words: Transition phase, Calcium Homeostasis, Small Ruminant, Maternal Nutrition Nonstandard abbreviations: bAP = bone-specific alkaline phosphatase activity; BMD = bone mineral density; calcitriol = 1,25-(OH)_2_ vitamin D; CTX = crosslaps;; ICTP = crosslinked carboxyterminal teleopeptide of type I collagen; VDR = vitamin D receptor

## Introduction

To avoid a drop in circulatory Ca, the organism has 3 options to stabilize the system: increasing absorption capacity at the gut barrier, limiting endogenous losses via the gastrointestinal tract and kidneys, and mobilizing Ca from the skeleton (Wilkens and Muscher-Banse, 2020). With respect to the regulation of Ca homeostasis, the animal species under study is a key factor to consider. For example, sheep and goats have been demonstrated to respond quite differently to Ca-restricted feeding, with sheep tending to mobilize bone stores more quickly, whereas goats attempt to compensate through more efficient absorption from the small intestinal mucosa (Wilkens et al., 2012a). A notable regulation of renal Ca excretion and reabsorption in response to changes in endogenous Ca status has been reported in mice (Hoenderop et al., 2002, van Cromphaut et al., 2003, Ko et al., 2009), but this mechanism has not been confirmed in ruminants, including sheep, goats (Herm et al., 2015), and cows (Wild et al., 2021). Adaptions to fluctuations in endogenous Ca levels are regulated by a complex system of endocrine factors that control the regulation of absorption, excretion, and bone remodeling. A detailed discussion of this system exceeds the scope of this introduction (see the reviews by Martín-Tereso and Martens (2014) and Wilkens and Muscher-Banse (2020)).

In lactating dairy ruminants, a sufficient endogenous Ca status is particularly important since milk supplies substantial amounts of Ca to offspring daily. Modern high-yielding dairy genotypes experience disproportionately increased Ca demands due to high milk production, despite the offspring’s Ca needs remaining relatively stable before and after parturition (Hernández-Castellano et al., 2020). In this context, the final weeks before parturition are critical. In high-yielding dairy cows, the sudden onset of lactation combined with reduced feed intake around parturition creates a risk of milk fever if bone mobilization is insufficient to compensate for dietary Ca deficits. This condition is typically accompanied by hypocalcemia, often co-occurring with hypophosphatemia and hypomagnesemia (Liesegang et al., 2007, Goff, 2008, Martín-Tereso and Martens, 2014). Symptoms manifest within 48 h postpartum and may include listlessness, coma, and sternal recumbence. The situation for sheep and goats is somewhat different. Due to their lower milk yield, meat sheep breeds rarely develop postpartum milk fever, though carrying twins or triplets increases the risk of antepartum hypocalcemia due to increased fetal Ca requirements during the final 4 weeks of gestation, when fetal skeleton calcification peaks. Sheep affected by hypocalcemia typically exhibit anorexia and stiff gait before progressing to sternal recumbence. Dairy sheep breeds and most goat breeds combine high milk yield potential with a greater likelihood of multiple pregnancies, making clinical cases of hypocalcemia likely both before and after parturition. High-yielding dairy sheep and goats develop a milk fever syndrome comparable to dairy cows but with a delayed onset, occurring 1-3 weeks postpartum. If antepartum symptoms arise, they tend to be less severe than those observed postpartum (Oetzel, 1988).

Several dietary and metabolic factors influence milk fever incidence, with most research focusing on dairy cows. Among these, DCAD, Mg, P, and Ca levels in antepartum diets have been widely studied (Lean et al., 2006). There has been ongoing debate regarding the optimal antepartum dietary Ca concentration, with some studies suggesting that Ca levels of 1-1.5% or even 2-3% in DM do not significantly impact milk fever incidence, while others recommend keeping dietary Ca below 1% in DM. An earlier meta-analysis attempted to clarify these inconsistencies. The authors developed 2 models using data from 51 trials, updating earlier findings from Oetzel (1991). They concluded that dietary Ca has a non-linear relationship with milk fever incidence, with both very low and very high concentrations appearing protective. Model I included antepartum DCAD, dietary Ca, P, Mg, and diet exposure time, while Model II replaced DCAD with antepartum dietary K and S. Both models suggested a quadratic effect of dietary Ca on milk fever pathogenesis (odds ratios: 0.131 and 0.115, respectively). Simulations with Model I, which varied dietary Ca from 0 to 2.5 % in DM while keeping all other variables stable at low-risk levels, produced a bell curve indicating that milk fever incidence peaked at ∼+3.8% when dietary Ca reached 1.35 % in DM. Based on this simulation, it was suggested that dietary Ca concentrations between 1.1-1.5% in DM should be avoided, whereas both lower and higher levels progressively reduce milk fever risk (Lean et al., 2006). These findings may explain inconsistencies across studies examining the interaction between dietary Ca and milk fever incidence. This raises an important question regarding whether similar effects occur in other livestock ruminants.

Studies on high dietary Ca levels in gestating sheep and goats are limited, and specific guidelines for preventing milk fever in these species are lacking – particularly in Switzerland. Our experimental design was informed by Lean et al. (2006), who found that antepartum dietary Ca levels between 1.1-1.5% in DM (compared to 0.6% in DM) could increase milk fever incidence by up to 3.8% in cows. This study aimed to evaluate the effects of high dietary Ca in dairy sheep and goats during the final 21 days of gestation on Ca homeostasis and bone metabolism while also providing preliminary data on potential risks extending into lactation. Given the naturally high environmental Ca concentrations in alpine regions such as Switzerland – where average forage Ca concentrations can reach 7 g/kg DM (Schlegel et al., 2016) – dietary Ca oversupply is not unlikely. Therefore, gestating sheep and goats from our in-house dairy herd were fed diets containing either 0.6% or 1.3% Ca in DM during the final 3 weeks antepartum, providing 1.15-fold and 2.54-fold the Swiss daily Ca intake recommendations (Agroscope, 2017). These levels were chosen to assess whether Ca intake below 1.1% in DM reduces milk fever risk while also testing an intake level approximating the hypocalcemia risk peak suggested by Lean et al. (2006). At the same time, Ca deficiency was deliberately avoided, as it would not have contributed to our research objectives. Observations continued until 56 days postpartum, during which animals were fed standard lactation diets containing recommended Ca levels based on Swiss guidelines (Agroscope, 2017). We hypothesized that sheep and goats consuming the higher Ca diet antepartum (equivalent to 30.5 g/day) would show signs of hypocalcemia at some point during the observation period, unlike those on the lower Ca diet. This design also allowed for comparative analysis of mineral and bone metabolism responses to dietary Ca overload in 2 species with distinct homeostatic strategies for managing Ca fluctuations (Wilkens et al., 2012b).

## Material and Methods

### Animals and Housing

This study was conducted at the research and laboratory facilities of the Institute of Animal Nutrition and Dietetics, Vetsuisse-Faculty, University of Zurich, Switzerland. A total of 10 and 12 multiparous gestating dairy sheep and goats (East Friesian Dairy Sheep, Saanen Goats), respectively, aged between 4 and 6 years were housed in individual pens with concrete floor and wood shavings as bedding material, from 4 weeks before the estimated parturition date. The imbalance in animal numbers between species was due to miscarriages of 2 sheep before the onset of the study. Both species were housed in separate rooms. The goats and sheep were housed individually until their delivery date, and afterwards, with their respective lambs and kids in lambing/kidding pens. The average body weight just before birth of goats and sheep was 95.2 ± SD1.5 kg and 101 ± SD2.22 kg, respectively, and 78.3 ± SD2.22 kg and 81.0 ± SD2.31 kg directly after birth, respectively.

### Diets and Experimental Design

Animals were randomly divided in 2 experimental groups according to their ear tag numbers using a random sequence generator (https://www.random.org/sequences/). The effective sample size for sheep and goats was 5 and 6 per group, respectively. The feeding trial started 3 weeks before the estimated day of parturition. The diets were adapted to the specific Swiss recommendations for energy and nutrient intake during gestation and lactation for sheep and goats (Agroscope, 2017). The 0.6 % in DM Ca groups received a balanced diet provided by grass hay (2^nd^ cut) and a commercial high energy concentrate (UFA 765, Herzogenbuchsee, Switzerland). Additionally, for the 1.3 % in DM Ca groups until parturition, the same concentrate was applied but mixed daily with 2.5 times (16.8 g Ca /day) the recommended daily Ca supply from CaCO_3_ (Table 1). From parturition onwards, all animals received lactation feeding to meet current Swiss feeding recommendations which accounted for 17.6 and 19.0 g Ca per day for sheep and goats, respectively (Table 1) (Agroscope, 2017).

**Table 1.**
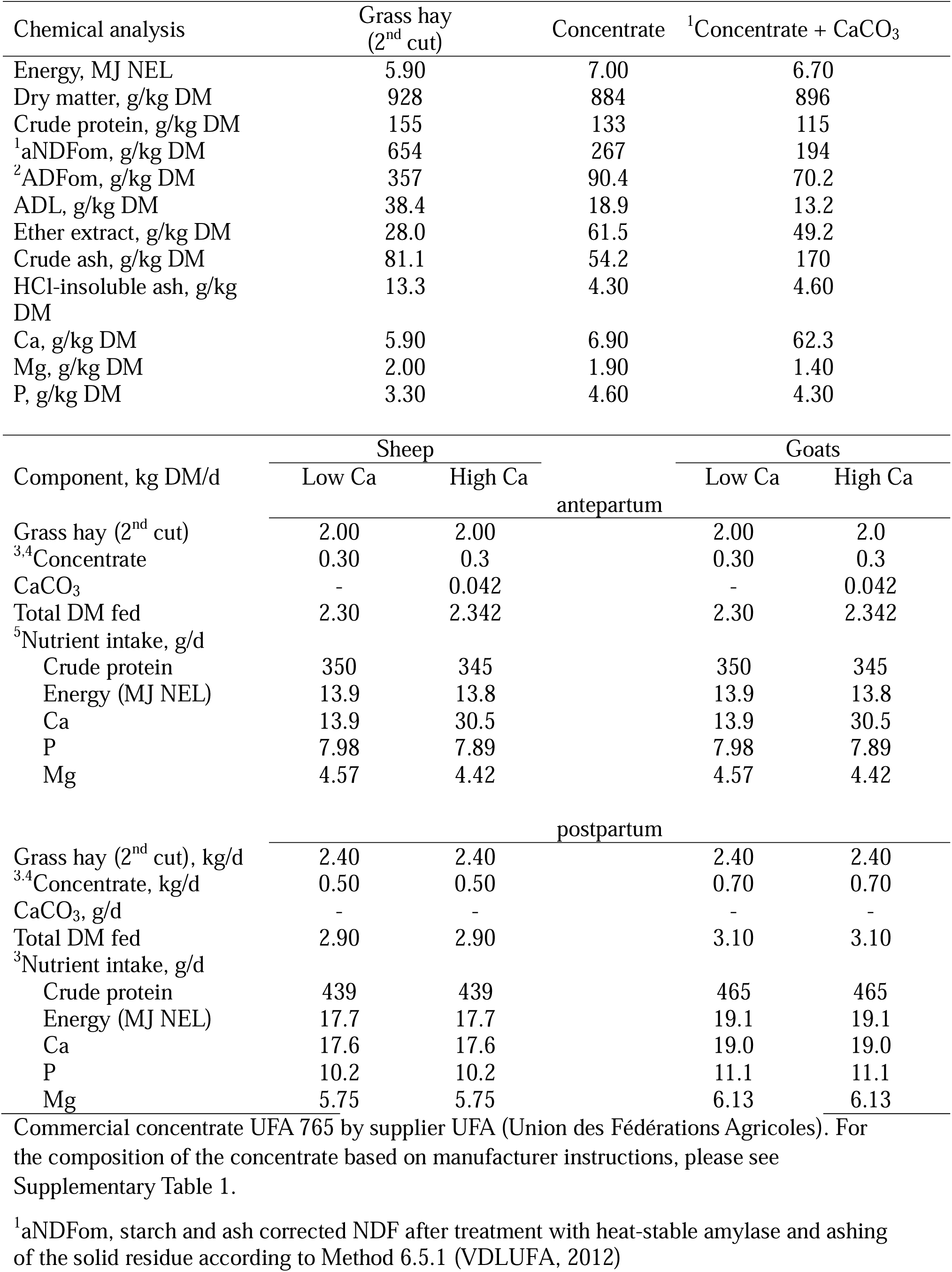
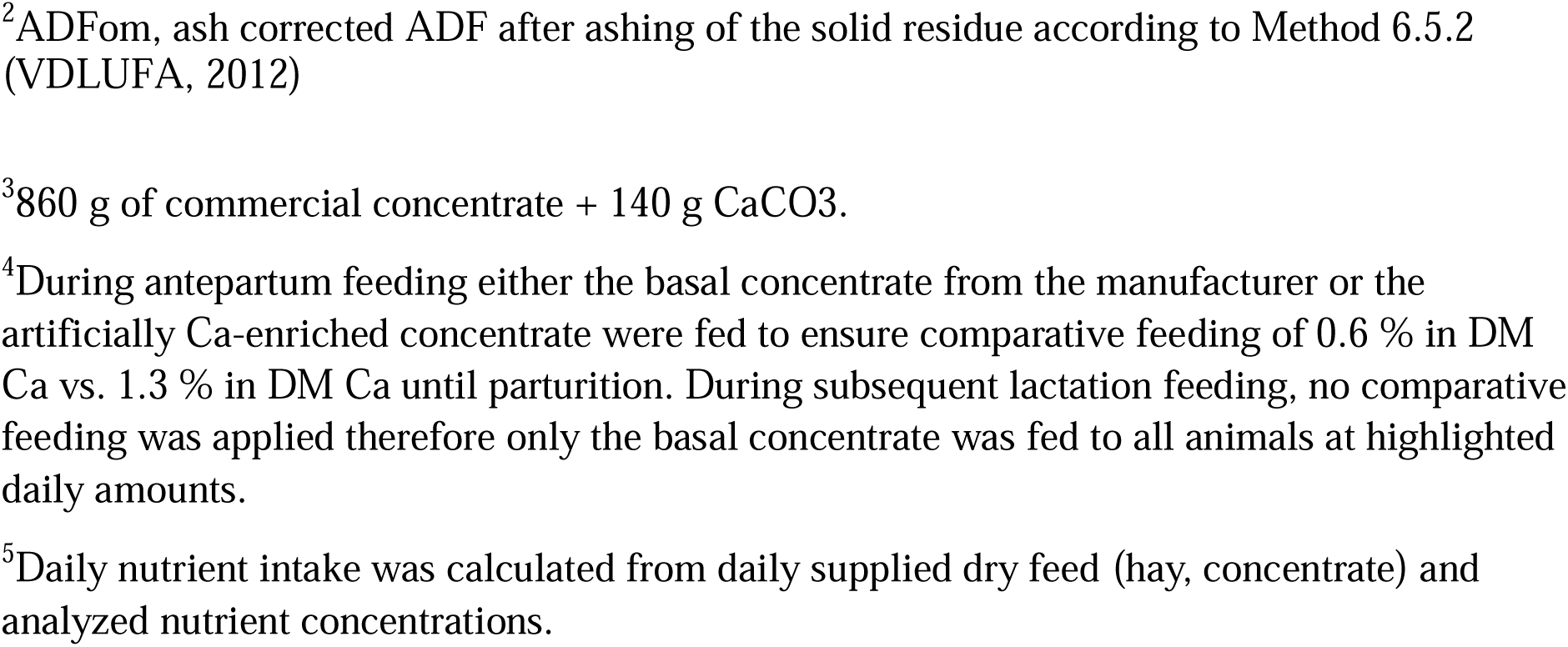
Chemical composition and daily intake levels of feedstuffs fed during the last 21 days antepartum until 56 days of lactation of dairy sheep and goats.

Throughout the study, animals were fed restrictive amounts of hay twice per day (2 kg/animal*d^-1^ antepartum and 2.4 kg/animal*d^-1^ postpartum for both species) and concentrate (0.3 kg/animal*d^-1^ antepartum for both species and 0.50 and 0.70 kg/animal*d^-1^ postpartum for sheep and goats, respectively) adapted to recommendations for antepartum and postpartum feeding (with the exception of Ca antepartum), and had ad libitum access to drinking water (Table 1). The restrictive feeding approach allowed for better feed intake control during single housing of animals. The composition of the commercial concentrate (UFA 765) as communicated by the manufacturer UFA (Union des Fédération Agricoles), is shown in Supplementary Table 1.

### Collection and handling of samples and growth parameters

Colostrum was hand milked from each animal immediately after parturition, and yield ranged from 0.5 to 1.2 L per animal in both species. After sampling 50 mL for subsequent analysis, colostrum was distributed equally over nursing bottles and supplied to the neonates.

Sheep and goats were weighed before morning feeding the day before and directly after parturition. Due to the suckling of the offspring, daily milk yield could not be assessed. The lambs and kids were weighed directly after parturition and once between 33-59 days postpartum, to determine the average daily weight gain during the suckling phase. The variation in the time interval between the 1^st^ and 2^nd^ weighing was due to lambs and kids being weaned at different ages for a different research purpose. To account for this variation in observational periods, we included the time interval between the 1^st^ weighing at birth and the 2^nd^ weighing (the age of each lamb and kid at 2^nd^ weighing) as a continuous variable in the statistical analysis of weight gain. This approach controls for potential age effects that could bias the estimation of treatment effects (see the subsection on statistical analysis below).

For this experiment, a closed batch of grass hay (2^nd^ cut) and concentrates was ordered and 3 representative samples were taken from each, following published standard methods 1.3 (“Regulations of the German Agricultural Society on the Sampling of Feed and Sample Treatment”) and 1.5 (“Sampling of Hay”) (VDLUFA, 2012) in order to investigate the feed value by chemical analysis. Feed samples were immediately subject to chemical analysis after sample collection.

Blood was collected daily for the 28 days before the estimated parturition and on days 1, 2, 3, 4, 5, 6, 7, 14, 28, and 56 postpartum, in all cases before morning feeding at 8 a.m. in serum monovettes (9 and 6 mL, Vacuette®, Greiner Bio-One Vacuette Schweiz GmbH) and centrifuged for serum collection (1580 x g, 10 min) after allowing coagulation at room temperature. The serum was stored at −80°C until further usage. After the actual parturition, samples of days −21, −4, −3, −2, −1, 0 (parturition), 1, 2, 3, 4, 5, 6, 7, 14, 28, and 56 were reserved for further analysis.

Urine was collected in a beaker upon spontaneous urination, in parallel to the blood sampling antepartum and on days 7 and 14 postpartum and homogenized to be separated into 2 equal subsamples. The 1st of the subsamples was acidified (12 mol/L HCl, pH 1-2) for later Ca determination, the other was subject to the later determination of creatinine. Both subsamples were stored at −20°C. After parturition, samples of days −21, −14, −7, 0, 7, and 14 were reserved for further usage. Fecal grab samples of each animal were collected in parallel to antepartum blood sampling, and samples collected on days −21, −2, −1, and 0 were subjected to further analysis. Samples were stored at −20°C until further usage.

All goats underwent colonic mucosal biopsies by veterinary personnel on days −21, −7, 0, and 7. An endoscope (PCF-20, Olympus) equipped with biopsy forceps was inserted 40 cm into the large intestine via the anus to collect 4 samples (1.5-2.0 mm diameter) per animal*time^-1^; 2 of these samples were immediately placed into 4% neutral buffered formalin and the remaining 2 samples were snap frozen in liquid nitrogen and stored at −80°C.

### Feed analysis

Feed was analyzed according to VDLUFA (2012) with the following methods. Dry matter was determined according to Method 3.1 (“Moisture, Water”) by oven drying at 105°C for 4 h. Net energy for lactation (Method 25.1 “Determination of Gas Production according to the Hohenheim Gas Test”) was assessed by quantification of cumulative gas production from rumen fluid over 24h at 39°C and regression estimation of NEL considering cumulative gas production and analyzed crude nutrients and fiber fractions (Society of Nutrition Physiology (GfE), 2008a, b). Crude protein determination (Method 4.1.1 “Determination of Crude Protein”) involved the Kjeldahl method of H_2_SO_4_ extraction in the presence of a catalyst, alkalization with NaOH, titration of NH_3_ and multiplying the assessed nitrogen concentration with a factor of 6.25. Starch and ash corrected neutral detergent fiber determination (aNDFom) (Method 6.5.1 “Determination of Neutral-Detergent-Fiber after amylase digestion (aNDF) as well as after amylase digestion and ashing (aNDFom)”) involved treatment with heat-stable amylase for 1h, subsequent cooking in neutral detergent solution, and ashing of the solid residue. Ash-corrected acid detergent fiber (ADFom) (Method 6.5.2 “Determination of acid-detergent fiber (ADF) and of acid detergent fiber after ashing (ADFom)”) was assessed by 1 h cooking in acid detergent solution and ashing of the solid residue. Acid Detergent Lignin (Method 6.5.3 “Determination of Acid Detergent Lignin (ADL)”) was quantified by incubating the starch-corrected ADF residue for 3 h at room temperature in 72% H_2_SO_4_ and subsequent ashing of the residue. The determination of the ether extract (Method 5.1.2 “Determination of Crude Fat”) involved the pre-extraction in hot HCl and subsequent extraction of the dry residue in petroleum ether. Crude ash (Method 8.1 “Determination of Crude Ash”) was measured by dry ashing at 550 °C for 4 h, and HCl-insoluble ash (Method 8.2 “Determination of Hydrochloric Acid-insoluble Ash”) was determined by adding an aliquot of the crude ash residue to boiling HCl. The dietary mineral profile (Ca, P, Mg) (Method 10 “Determination of selected elements in plant material and feedstuffs with ICP-OES”) involved the sample ashing in a furnace, extraction with HCl, with a deviation from the protocol, by measuring the solutions not via ICP-OES but by flame atomic absorption spectrometry (flame-AAS) (ContrAA 700, Analytik Jena).

### Analysis of blood serum parameters of bone and mineral homeostasis

Blood serum was analyzed for osteocalcin, 1,25-(OH)_2_ vitamin D (**calcitriol**), bone-specific alkaline phosphatase (**bAP**), crosslinked carboxyterminal teleopeptide of type I collagen (**ICTP**), and crosslaps (**CTX**).

Serum osteocalcin was quantified with a commercial EIA test kit (MikroVue Osteocalcin EIA Kit; Quidel Corporation). Intra- and interassay CVs were 5.00% and 5.75% with a sensitivity of 0.45 ng/mL. Serum calcitriol determination was conducted with an RIA kit (1,25VitD RIA, sheep-anti 1,25VitD antibody; Immunodiagnostics Systems GmbH). Intra- and interassay CVs were 9.0% and 9.3%, respectively, with a sensitivity of 8 pmol/L. Serum bAP was analyzed with an immunoassay via a monoclonal anti-bAP and para-nitrophenylphosphate as a substrate (Alkphase-BTM^TM^; Metra Biosystems). Intra- and interassay CVs were 0.58 and 0.52.

Serum ICTP was measured by a RIA immunoassay (UniQ® ICTP RIA, DiaSource) with intra- and interassay CVs of 5.6% and 10.0%, respectively, and a sensitivity of 0.6 µg/L. Serum CTX was assayed by a commercial ELISA assay (Serum CrossLaps (CTX-I) ELISA; Immunodiagnostics Systems GmbH) with intra- and interassay CVs of 1.7% and 2.5% at a sensitivity of 0.02 ng/mL.

For CTX, ICTP, and calcitriol, data gaps at −4 days and for CTX additionally for −21 days, as depicted in Figures 5 and 6, respectively, were due to compromised sub-samples. All other sampling decisions were intentional, based on a balance between data relevance, workload, and monetary considerations.

### Analysis of Ca and creatinine in biological matrices

Serum Ca was measured via flame-AAS after microwave wet digestion of 2 mL of serum per technical replicate in 65% HNO_3_ and 30% H_2_O_2_ (Ethos, MLS GmbH) against a dilution gradient (0.2, 0.4, 0.6, 0.8, 1.0 mg/L) of a commercial AAS-certified standard (ROTI^®^Star 1000 mg/L Ca, in 2 % HNO_3_; Roth).

Dried Feces (105°C, 4 h) was milled (0.5 mm) and 500 mg per technical replicate was subjected to Ca analysis as described for serum above.

Colostrum was defatted by centrifugation (4000 x g, 4°C, 30 min), and was subjected to the same procedures for sample preparation and Ca analysis as described for feces and blood serum.

Urine was analyzed for total Ca with the method described for serum and further analyzed for creatinine by a kinetic color test using a commercial kit (Crea Jaffe DIA00540, Diatools AG) on the Cobas Mira autoanalyzer with an intra- and interassay CV of 0.83% and 0.85% at a sensitivity of 18 µmol/L. The urinary Ca excretion was expressed as ratio to urinary creatinine since the material could not be sampled quantitatively. Creatinine is generally excreted at a relatively constant rate, making it a useful reference compound to normalize mineral concentrations for differences in urine dilution caused by hydration status, collection timing or other individual factors (Sullivan et al., 2012).

### Vitamin D receptor immunoreactivity

Goat colonic mucosal biopsies were subjected to semi-quantitative immunohistochemical determination of vitamin D receptor (**VDR**) abundance as originally described by Liesegang et al. (2008) and Sidler-Lauff et al. (2010). Caprine colonic VDR was labelled after rinsing the tissue with TBS buffer and incubation for 80 min with a biotinylated rat monoclonal antibody (9A7γ, ThermoScientific) at a 1:200 dilution in Tris-buffered saline against unspiked Tris-buffered saline on an adjacent colonic section as negative control and duodenal cross sections of pigs (obtained from the in-house slaughtering facility of the Vetsuisse-Faculty, University of Zurich) (aa 89-105) as positive control based on earlier reported identical cross-reactivity (Milde et al., 1989). For sheep, no suitable antibody was available at the time of study.

Nuclear immunoreactivity was assessed with a binocular microscope (DMLB®, Leica) with a digital camera (DC480, Leica) (magnification 400x, 5-7 microscopic fields). For each sample, 500 cells of the following cell types were evaluated for nuclear staining intensities: surface epithelial cells, intermediate glandular/crypt epithelial cells, superficial glandular/crypt epithelial cells, basal glandular/crypt epithelial cells. Staining intensities of nuclei were scored 0, 0.5, 1, 2, 3 for negative, very weak, weak, intermediate, and strong, corresponding to the absence of brown staining (only blue counter staining), light brown, brown, and dark brown, respectively. From this an immune reactivity score was calculated with the following formula:

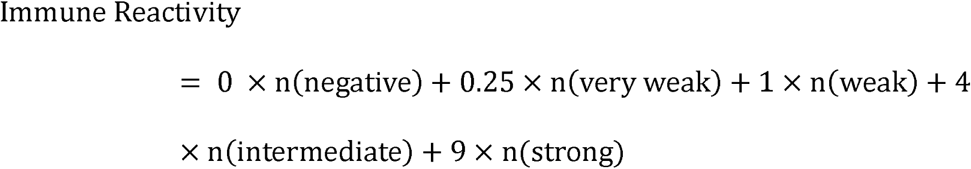

Where n is the number of cells within 500 cells expressing a respective staining intensity, with a range of calculated immune reactivity scores between 0 (no specific staining) to 4500 (all 500 cells with strong staining).

### Bone mineral density

Bone mineral density (**BMD**) was measured with peripheral quantitative computer tomography (Stratec XCT 2000 bone scanner, Stratec Medizinaltechnik GmbH) in the middle of the left metatarsus after length determination with a caliper. Measurements were taken in the mid-diaphysis and cortical BMD was automatically calculated (cortical mode 2, >640 mg/cm^3^ cortical bone threshold).

### Statistical analysis

The individual animal was the smallest experimental unit. All data analysis was performed with SAS 9.4 (SAS Institute Inc.) applying the procedure MIXED. Repeated measures mixed models were applied to analyze the effects of the fixed variables “diet Ca”, “species”, “time”, and interactions considering the individual animal nested under the respective feeding group as a random factor. Repeated measures mixed modelling involved in any case the mapping of covariance structures appropriate for the respective data structure following guidance by Wang and Goonewardene (2004). Due to the uneven spacing of times of observation for respective datasets, covariance structures like first-order autoregressive (AR(1)) and related structures (e.g., Toeplitz), which require an equal spacing of time, were not considered. The Bayesian Information Criterion (BIC) served as model fit statistic, and we chose the candidate yielding the lowest BIC. The respectively mapped covariance structure for different repeated measures datasets were as follows: compound symmetry (serum Ca, urinary Ca), heterogeneous compound symmetry (BMD), first-order ante-dependent (serum bAP, serum ICTP, serum CTX, serum calcitriol, serum osteocalcin, fecal Ca). Due to the limited availability of suitable antibodies, the VDR protein expression over time was only analyzed in colonic biopsies of goats, therefore, no species effect could be estimated.

The colostral Ca concentration of sheep and goats was analyzed with an endpoint mixed model with the fixed effect of “diet Ca”, “species”, and interaction (diet Ca*species), again including the individual animal nested under the respective feeding group as a random effect. Birth weights and daily weight gain of the offspring were also analyzed with an endpoint mixed model accounting for the fixed effects of “diet Ca”, “species”, “sex”, and interactions (diet Ca*species, diet Ca*sex) including the individual animal nested under the respective mother as a random effect. For the analysis of average daily weight gain, the animals’ age at the 2^nd^ weighing was included as a continuous control variable to account for potential age-related bias in estimating average daily weight gain. If the residuals of the respective mixed models did not meet the assumption of normality, the analysis was repeated after Log_10_-transformation of data and displayed as such. This applied to the following datasets: fecal Ca, urinary Ca, serum bAP, serum ICTP, serum CTX. Data were presented in any case as least square mean values with Tukey-adjusted 95% confidence limits (Cl[upper, lower]). *P* ≤ 0.05 was defined as the threshold for a Type I Error.

Earlier datasets were used for prospective power analyses during study planning. Therefore, effect sizes of ante and postpartum adaptation of assessed serum parameters were determined with the respective datasets of Liesegang et al. (2007). Effect sizes of VDR immune reactivity in the goat colon to dietary Ca variation were determined according to the respective dataset of Sidler-Lauff et al. (2010). The BMD effect sizes of their fluctuation from gestation to lactation of milk goats and sheep were estimated with the respective data of Liesegang et al. (2007). The effect sizes and minimum number of replicates required to achieve statistical significance (with a minimum statistical power of 1-β = 0.8 at α = 0.05) (McDonald, 2014) were estimated using R version 4.4.2 (The R Project for Statistical Computing). This analysis employed the SIMR package following the approach outlined by Green and MacLeod (2016), with the modification of using the previously published datasets mentioned earlier instead of simulated datasets. Based on these estimations, the anticipated minimum power of ≥0.8 was reached in any case assuming 6 replicates per feeding group, applying the models described above.

## Results

### Animal health and development

All sheep and goats remained healthy throughout the entire study period with no signs of pathology or hypocalcemia, based on continuous veterinary surveillance. Sheep showed 5 cases of triplets, 4 cases of twins and 1 single birth (n = 24 lambs in total) whereas most goats gave birth to twins with 1 case of triplets and 2 cases of single births (n = 23 kids in total). The sex ratio of sheep and goat offspring was (m/f) 0.71 and 1.30, respectively.

All lambs and kids were born with species-dependent birth weights (4.62 Cl[4.22, 5.01] vs. 5.15 Cl[4.80, 5.51] kg for kits vs lambs; *P* = 0.06) irrespective of sex (4.71 Cl[4.35, 5.07] vs. 5.06 Cl[4.71, 5.41] kg for female vs. male; *P* = 0.16) and were unaffected by the mothers’ dietary Ca supply antepartum (4.89 Cl[4.53, 5.23] vs. 4.88 Cl[4.54, 5.22] kg at 0.6% vs. 1.3% in DM, respectively; *P* = 0.94) (Figure 1A). However, lambs and kids of mothers fed high Ca antepartum showed significantly lower daily weight gain from milk intake than animals fed low Ca (0.18 Cl[0.17, 0.20] vs. 0.22 Cl[0.20, 0.24] kg/d at 1.3% vs. 0.6% in DM, respectively; *P* = 0.02) irrespective of sex and species, which expressed distinct effects on the animals’ growth performance (0.19 Cl[0.17, 0.21] vs. 0.21 Cl[0.19, 0.23] kg/d for female vs. male; *P* = 0.06; 0.25 [0.23, 0.27] vs. 0.15 [0.13, 0.17] kg/d for kids vs. lambs; *P* < 0.0001) (Figure 1B). Supplementary Table 2 highlights all LSmeans and associated Tukey-adjusted 95% confidence limits of birth weights and daily weight gain of offspring as related to species, sex, and dietary Ca supply of the mothers antepartum.

**Figure 1.**
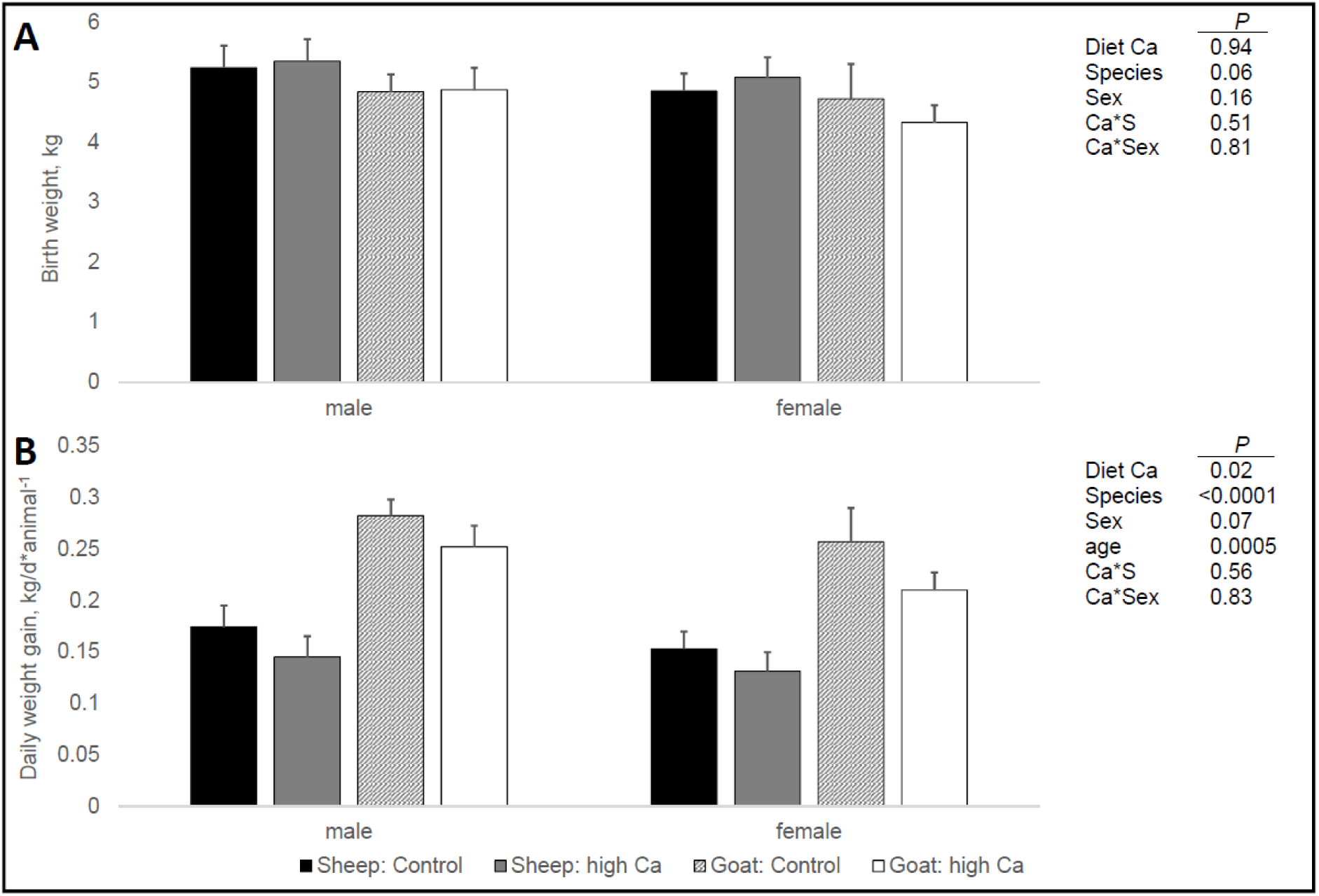
Effects of varying dietary Ca (0.6 vs. 1.3% in DM) of dairy sheep and goats during the last 3 weeks antepartum as well as sex and species of offspring on the birth weight (A) and daily weight gain during the suckling phase (B). Effective sample size was 5 sheep and 6 goats per group, which gave birth to 24 and 23 lambs and kids in total, respectively. Data is presented as least square mean values with standard errors based on linear mixed model analysis. “Age” refers to the time of 2^nd^ weighing during the suckling phase, which differed between individual animals due to management reasons (range: 33-59 days postpartum) and was therefore introduced as a continuous control variable to the statistical analysis of daily weight gain data. Ca*S and Ca*Sex refer to statistical interactions between diet Ca and species of animal or sex of offspring, respectively. *P* ≤ 0.05 was defined as statistical type I error threshold.

### Quantitative markers of calcium metabolism

Analyzed fecal Ca concentrations during the observation period ranged between 1.46 and 5.19 % in DM in sheep as well as 1.17 and 14.9 % in DM in goats whereas the ranges for urinary Ca excretion were 0.005 and 0.41 mmol/mmol creatinine in sheep as well as 0.004 and 1.55 mmol/mmol creatinine in goats. Fecal and urinary data had to be log_10_-transformed to meet the precondition of normality, accounting for fecal Ca ranges of 0.16 and 0.71 log_10_(% in DM) for sheep as well as 0.07 and 1.17 log_10_(% in DM) for goats. The respective log-transformed urinary Ca excretion ranges were −2.31 and −0.38 log_10_(mmol/mmol creatinine) in sheep as well as −2.42 and −0.19 log_10_(mmol/mmol creatinine) in goats. Supplementary Table 3 highlights all LSmeans and associated Tukey-adjusted 95% confidence limits of fecal Ca concentrations and urinary Ca excretion, expressed in the original measurement scale as related to species, time, and dietary Ca supply antepartum.

Higher supply with dietary Ca antepartum significantly increased fecal log_10_-Ca concentrations until parturition by 1.43-fold vs. 2.87-fold in sheep vs. goats and in any case significantly above the control level (0.21 Cl[0.07, 0.34] vs. 0.62 Cl[0.48, 0.75] and 0.35 Cl[0.22, 0.48] vs. 0.66 Cl[0.53, 0.79] log_10_(% in fecal DM) in 0.6% vs. 1.3% in DM of sheep and goats, respectively; *P* < 0.0001) (Figure 2A). On average, sheep showed lower fecal Ca than goats (0.41 Cl[0.36, 0.46] vs. 0.47 Cl[0.43, 0.52] log10(% in fecal DM) for sheep vs. goats; *P* = 0.05) but the response patterns over time were different. Sheep in the low Ca group slightly declined fecal Ca until parturition to 0.64-fold of the level at −21 days antepartum, while sheep in the high Ca group responded opposite to this as mentioned before (1.43-fold increase until parturition). In contrast, both control and high Ca goats increased the levels until parturition but in different magnitudes (1.75-fold vs. 2.87-fold for low vs. high Ca). This difference in time patterns between species was reflected by a strong statistical interaction between species and time (*P* < 0.0001).

**Figure 2.**
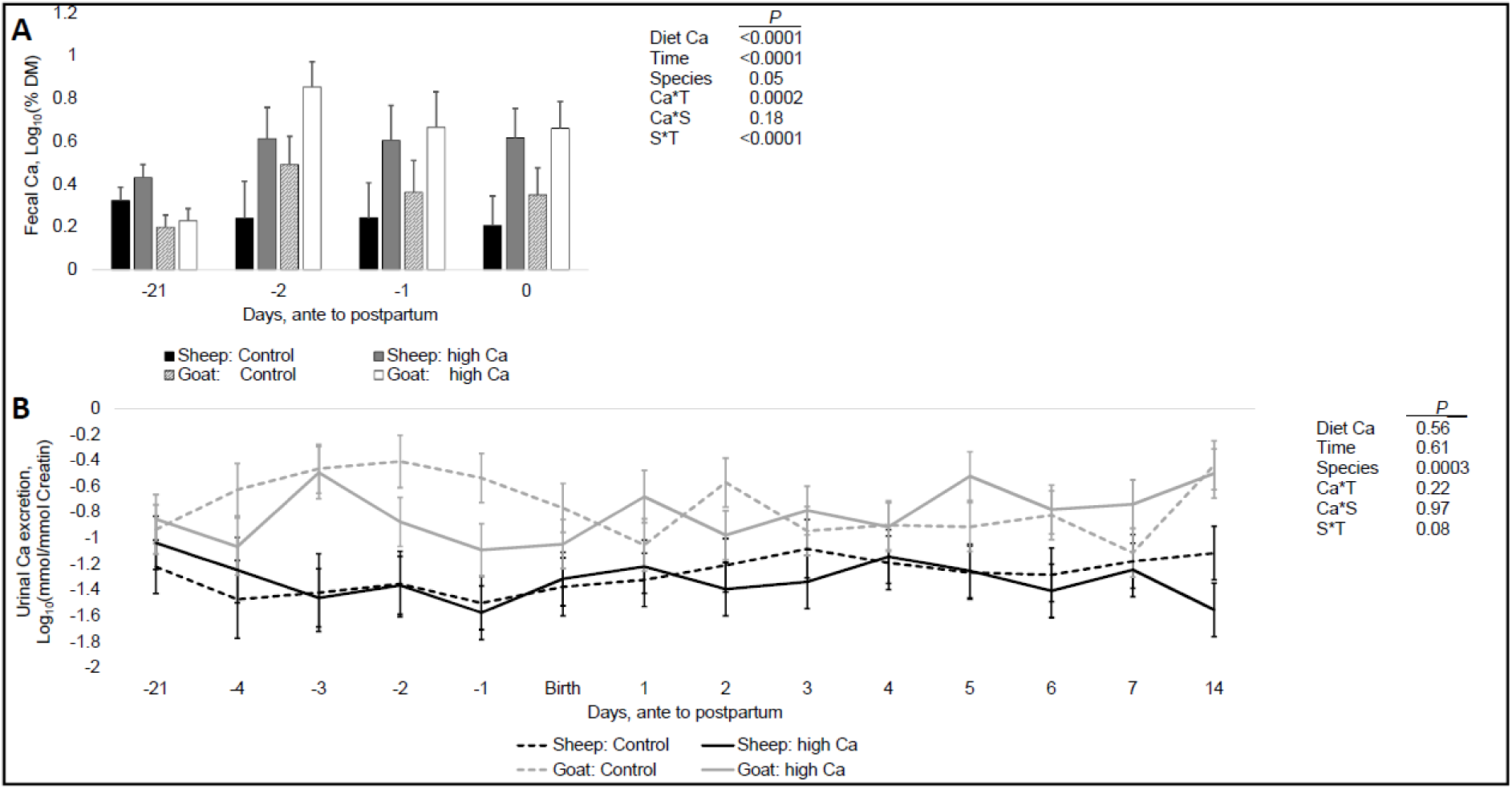
Effects of varying dietary Ca (0.6 vs. 1.3% in DM) of dairy sheep and goats during the last 3 weeks antepartum as well as time and animal species on the fecal Ca concentration (A) and urinary Ca excretion (B) from 3 weeks antepartum to parturition and 14 days postpartum, respectively. Effective sample size was 5 sheep and 6 goats per group. Data is presented as Log_10_-transformed least square mean values with standard errors, respectively, based on repeated measures linear mixed model analysis. Ca*T, Ca*S and S*T refer to statistical interactions between diet Ca and animal species or time, respectively, as well as animal species and time. *P* ≤ 0.05 was defined as statistical type I error threshold.

In contrast, the feeding regime had no effect on the urinary Ca excretion (−1.04 Cl[−1.22, - 0.86] vs. −0.96 Cl[−1.14, −0.79] log10(mmol/mmol creatinine) at 0.6% vs. 1.3% in DM; *P* = 0.56) (Figure 2B). Values fluctuated over time but a clear trend in adaptation of urinary losses was not evident (*P* = 0.61). The only significant effect was in comparison between sheep and goats irrespective of treatment or time, with goats excreting on average 3.26-fold the levels of sheep (−1.26 Cl[−1.44, −1.07] vs. −0.75 Cl[−0.91, −0.58] log_10_(mmol/mmol creatinine) in sheep vs. goats; *P* = 0.0003).

Figure 3 shows the response of serum Ca and colostral Ca to varying Ca feeding antepartum. Serum Ca was not affected by the antepartum feeding (2.47 Cl[2.35, 2.59] vs. 2.49 Cl[2.37, 2.61] mmol/L at 0.6% vs. 1.3% in DM; *P* = 0.80) and gradually and significantly (*P* < 0.0001) decreased independent of animal species towards the initial 4 days postpartum to gradually increase to previous levels during the further course of the observation period (Figure 3A). In any case, sheep showed on average 1.16-fold higher serum Ca levels than goats (2.29 Cl[2.17, 2.40] vs. 2.67 Cl[2.54, 2.79] mmol/L in goats vs. sheep; *P* = 0.0003).

**Figure 3.**
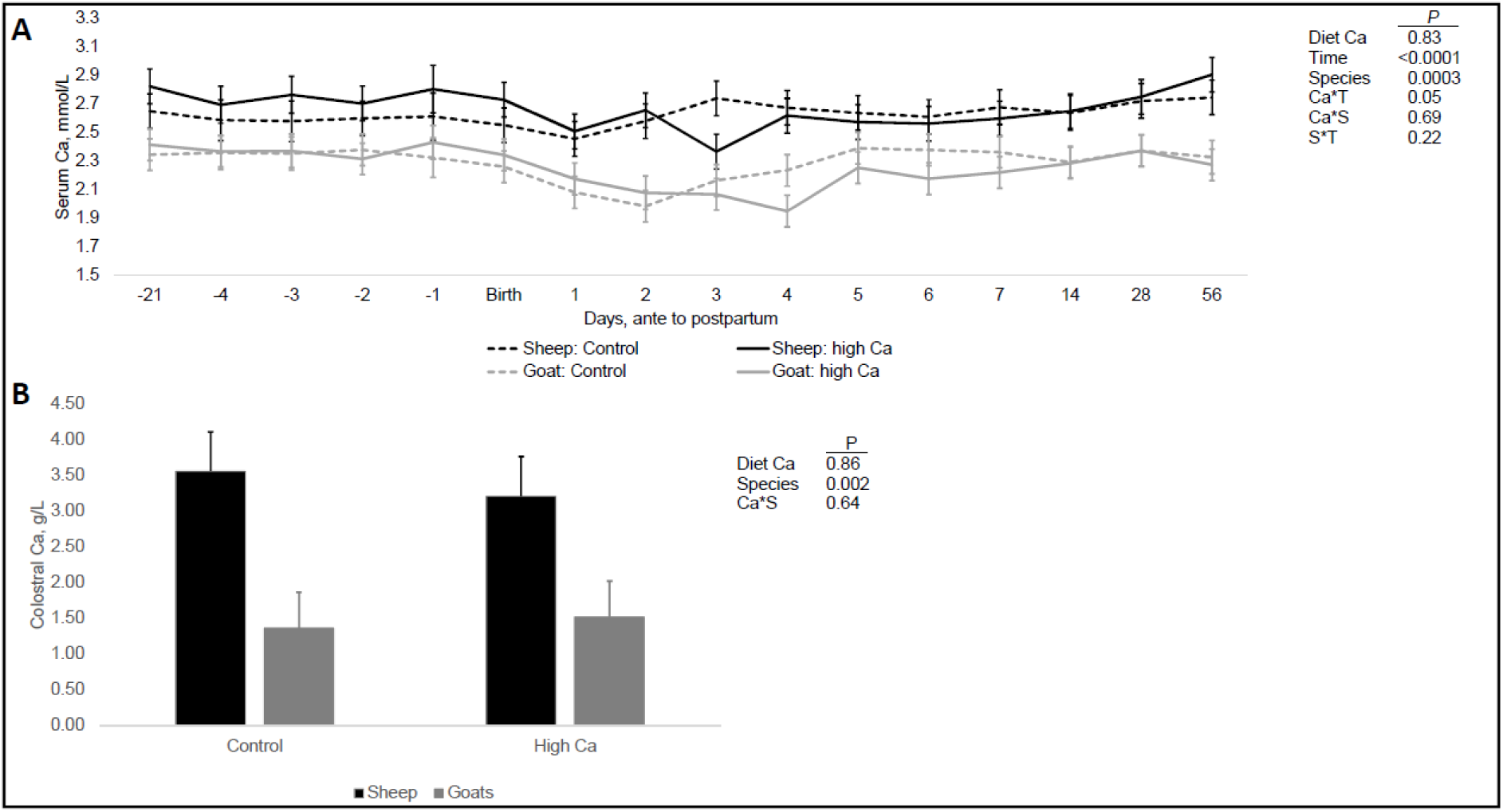
Effects of varying dietary Ca (0.6 vs. 1.3% in DM) of dairy sheep and goats during the last 3 weeks antepartum as well as time and animal species on serum Ca (A) and colostral Ca (B) from 3 weeks antepartum to parturition and 56 days postpartum, respectively. Effective sample size was 5 sheep and 6 goats per group. Data is presented as least square mean values with standard errors based on repeated measures linear mixed model analysis and regular linear mixed model analysis, respectively. Ca*T, Ca*S and S*T refer to statistical interactions between diet Ca and animal species or time, respectively, as well as animal species and time. *P* ≤ 0.05 was defined as statistical type I error threshold.

The colostral Ca concentration (Figure 3B) was 2.35-fold higher in sheep than in goats (1.44 Cl[0.69, 2.19] vs. 3.38 Cl[2.56, 4.20] g/L in goats vs. sheep; *P* = 0.002) but was not affected by the antepartum Ca feeding regime (2.36 Cl[1.58, 3.14] vs. 2.46 Cl[1.67, 3.24] g/L at 0.6% vs 1.3% Ca in DM; *P* = 0.86) (Figure 3B).

Supplementary Table 4 highlights all LSmeans and associated Tukey-adjusted 95% confidence limits of serum Ca and colostral Ca as related to species, time, and dietary Ca supply antepartum.

Figure 4 highlights the BMD of dairy sheep and dairy goats from −14 day antepartum to 56 days postpartum, as affected by Ca feeding during the last 3 weeks antepartum, time and animal species. Higher Ca feeding antepartum did not affect BMD (868 Cl[845, 891] vs. 877 Cl[854, 900] mg/cm^3^ at 0.6% vs. 1.3% in DM; *P* = 0.63), which significantly declined towards parturition from 932 Cl[904, 960] to 864 Cl[807, 922] mg/cm^3^ in sheep and 845 Cl[817, 873] to 782 Cl[738, 826] mg/cm^3^ in goats, to gradually increase postpartum until day 56 of lactation where it roughly approached in both species the initial level at the beginning of the observation (Sheep: 944 Cl[920, 967] mg/cm^3^, Goats: 846 Cl[823, 868] mg/cm^3^ *P* < 0.0001). On average, sheep showed 1.08fold higher BMD than goats (839 Cl[808, 853] vs. 913 Cl[888, 938] mg/cm^3^ in goats vs. sheep; *P* < 0.0001).

**Figure 4.**
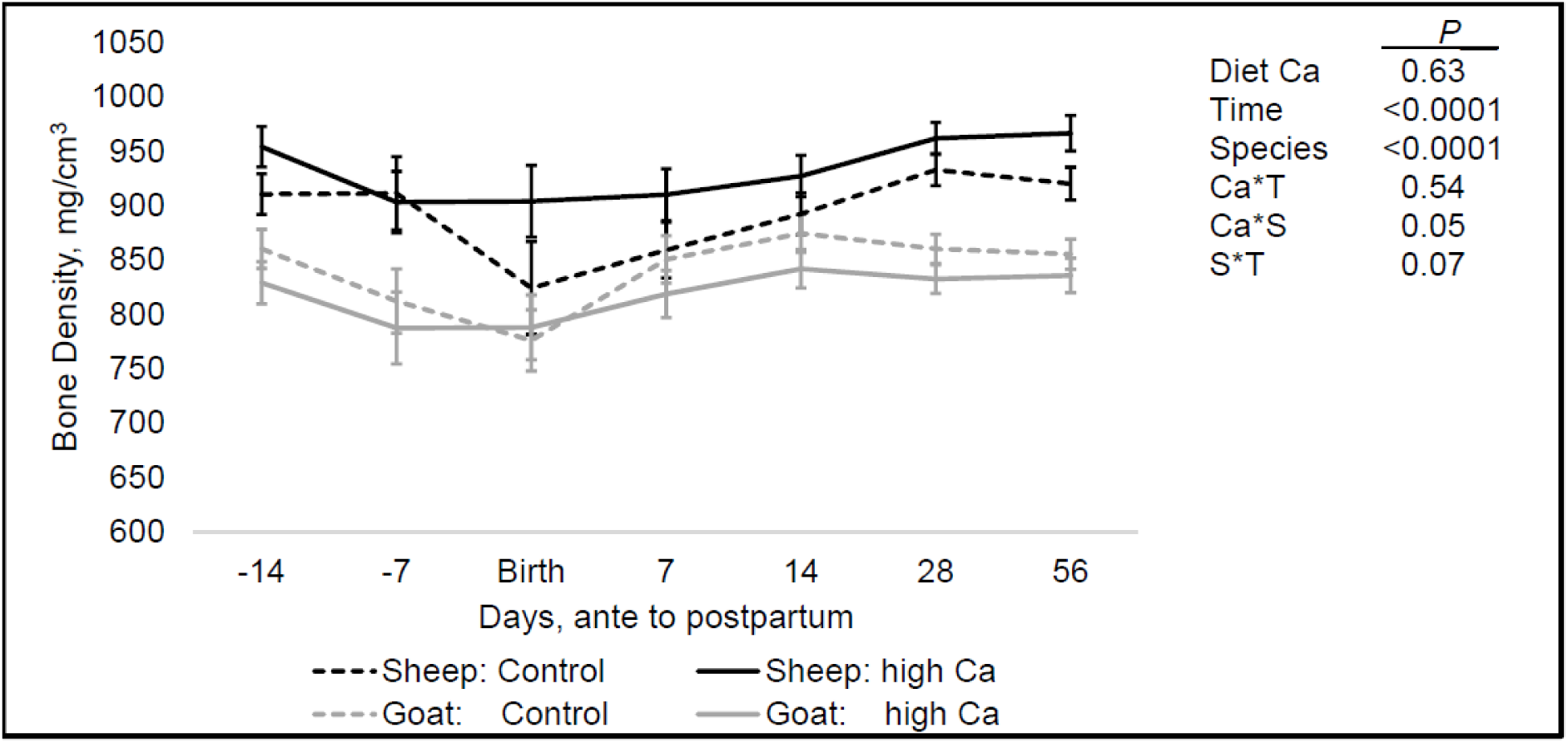
Effects of varying dietary Ca (0.6 vs. 1.3% in DM) of dairy sheep and goats during the last 3 weeks antepartum as well as time and animal species on BMD from 2 weeks antepartum to 56 days postpartum. Effective sample size was 5 sheep and 6 goats per group. Data is presented as least square mean values with standard errors based on repeated measures linear mixed model analysis. Ca*T, Ca*S and S*T refer to statistical interactions between diet Ca and animal species or time, respectively, as well as animal species and time. *P* ≤ 0.05 was defined as statistical type I error threshold.

Supplementary Table 5 highlights all LSmeans and associated Tukey-adjusted 95% confidence limits of BMD as related to species, time, and dietary Ca supply antepartum.

### Serum calcitriol and markers of bone formation

Serum calcitriol, bAP, and osteocalcin of sheep and goats over time in response to varying Ca feeding antepartum are shown in Figure 5. Serum calcitriol was 1.98-fold higher in goats than in sheep (70.2 Cl[56.8, 83.6] vs. 139 Cl[128, 149] pmol/L in sheep vs. goats; *P* < 0.0001) but not different between antepartum feeding groups (103 Cl[90.1, 117] vs. 105 Cl[92.3, 119] pmol/L at 0.6% vs. 1.3% in DM; *P* = 0.75) (Figure 5A). Serum levels significantly changed over time (*P* < 0.0001) which interacted with species (*P* < 0.0001) as follows. In sheep, serum levels reduced towards parturition and fluctuated throughout, whereas in goats serum levels increased until 4 days postpartum, then turned around and approached initial levels until 56 days postpartum.

**Figure 5.**
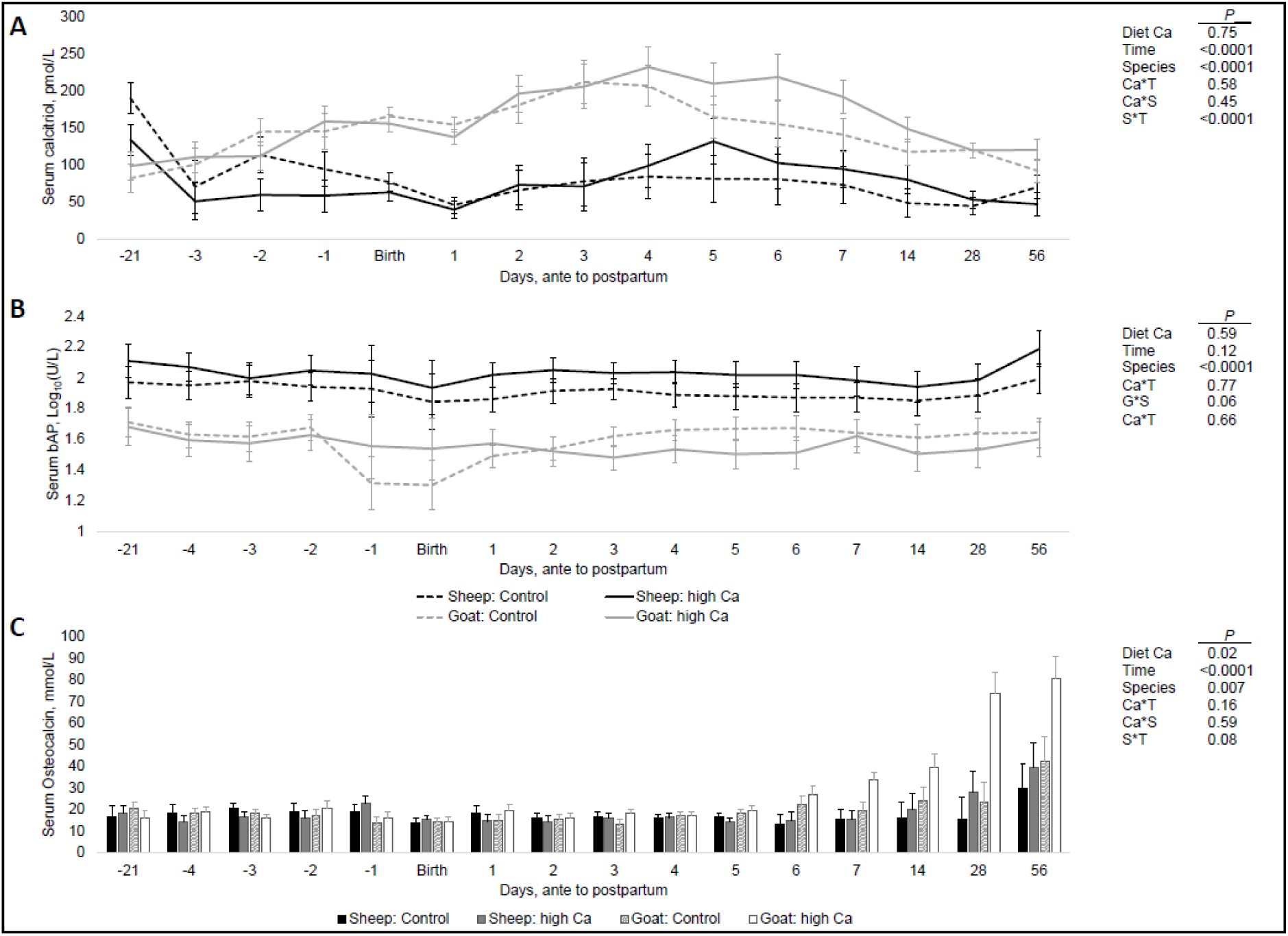
Effects of varying dietary Ca (0.6 vs. 1.3% in DM) of dairy sheep and goats during the last 3 weeks antepartum as well as time and animal species on serum calcitriol (A), serum bAP (B), and serum osteocalcin (C) from 3 weeks and 5 days antepartum, respectively, to 56 days postpartum. Effective sample size was 5 sheep and 6 goats per group. Data is presented as least square mean values (Log_10_-scale in case of bAP activity) with standard errors based on repeated measures linear mixed model analysis. Ca*T, Ca*S and S*T refer to statistical interactions between diet Ca and animal species or time, respectively, as well as animal species and time. *P* ≤ 0.05 was defined as statistical type I error threshold. Due to compromised sub-samples for analysis of calcitriol at −4 days antepartum, we cannot provide the respective data.

Serum activity of bAP ranged between 38 and 251 U/L in sheep as well as 21 and 117 U/L in goats. After Log_10_-transformation to meet the precondition of normality, these ranges were 1.58 and 2.40 Log_10_(U/L) in sheep as well as 1.32 and 2.07 Log_10_(U/L) in goats. Feeding group (1.76 Cl[1.67, 1.85] vs. 1.80 Cl[1.70, 1.90] log_10_(U/L) at 0.6% vs. 1.3% in DM; *P* = 0.59) expressed no significant effect on bAP (Figure 5B). This was also true for time, though control goats showed a transient numerical drop by ∼22% of serum activity around parturition (*P* = 0.12). Sheep showed on average 1.25-fold higher serum activities than goats (1.58 Cl[1.49, 1.68] vs. 1.97 Cl[1.888, 2.07] log_10_(U/L) in goats vs. sheep; *P* < 0.0001).

Serum osteocalcin remained stable and comparable between species from −5 days antepartum to 5 days and 7 days postpartum, respectively, after which it gradually increased on average by 2.27-fold in sheep and 3.23-fold in goats until 56 days postpartum (*P* < 0.0001) (Figure 5C). This difference in magnitude was significant between species (*P* = 0.007). Most notably, this increase in serum osteocalcin was significantly higher in those sheep and especially goats who received higher dietary Ca during the last 3 weeks antepartum (29.9 Cl[5.62, 54.1] vs. 39.6 Cl[15.3, 63.8] and 42.2 Cl[17.8, 66.6] vs. 80.5 Cl[58.4, 103] mmol/L at 0.6% vs. 1.3% in DM in sheep and goats, respectively; *P* = 0.02).

Supplementary Table 6 highlights all LSmeans and associated Tukey-adjusted 95% confidence limits of serum calcitriol, bAP, and osteocalcin, expressed in the original measurement scale, also in the case of bAP, as related to species, time, and dietary Ca supply antepartum.

### Serum markers of bone resorption

Serum concentrations of ICTP ranged between 4.87 and 90.2 µg/L in sheep as well as 5.75 and 57.6 µg/L in goats, while CTX ranged between 0.50 and 10.3 ng/mL in sheep as well as 0.35 and 8.72 ng/mL in goats. After Log_10_-transformation to meet the precondition of normality, these ranges were 0.69 and 1.95 log_10_(µg/L) as well as −0.30 and 1.01 log_10_(µg/L) for ICTP, and 0.76 and 1.76 log_10_(ng/mL) as well as −0.46 and 0.94 log_10_(ng/mL) for CTX. Supplementary Table 7 highlights all LSmeans and associated Tukey-adjusted 95% confidence limits of serum ICTP and CTX, expressed in the original measurement scale, as related to species, time, and dietary Ca supply antepartum. Serum ICTP showed no marked changes in both species until parturition, afterwards it significantly increased over the initial 2 days postpartum (1.49fold vs. 1.60fold in sheep vs. goats) and held a plateau until 7 days postpartum from which on the levels gradually reduced over the following weeks to approach initial values at 56 days postpartum (*P* < 0.0001) (Figure 6A). Average serum levels were significantly higher for sheep than goats (1.19 Cl[1.14, 1.23] vs. 1.30 Cl[1.26, 1.35] log_10_(µg/L) in goats vs. sheep; *P* < 0.0001) with a significant interaction between species and time reflecting fluctuations in differences between species over given times (*P* = 0.001). The feeding antepartum did not express any significant effect on serum ICTP levels of dairy sheep and goats (1.24 Cl[1.20, 1.29] vs. 1.25 Cl[1.20, 1.29] log_10_(µg/L) at 1.3% vs. 0.6% in DM; *P* = 0.98).

**Figure 6.**
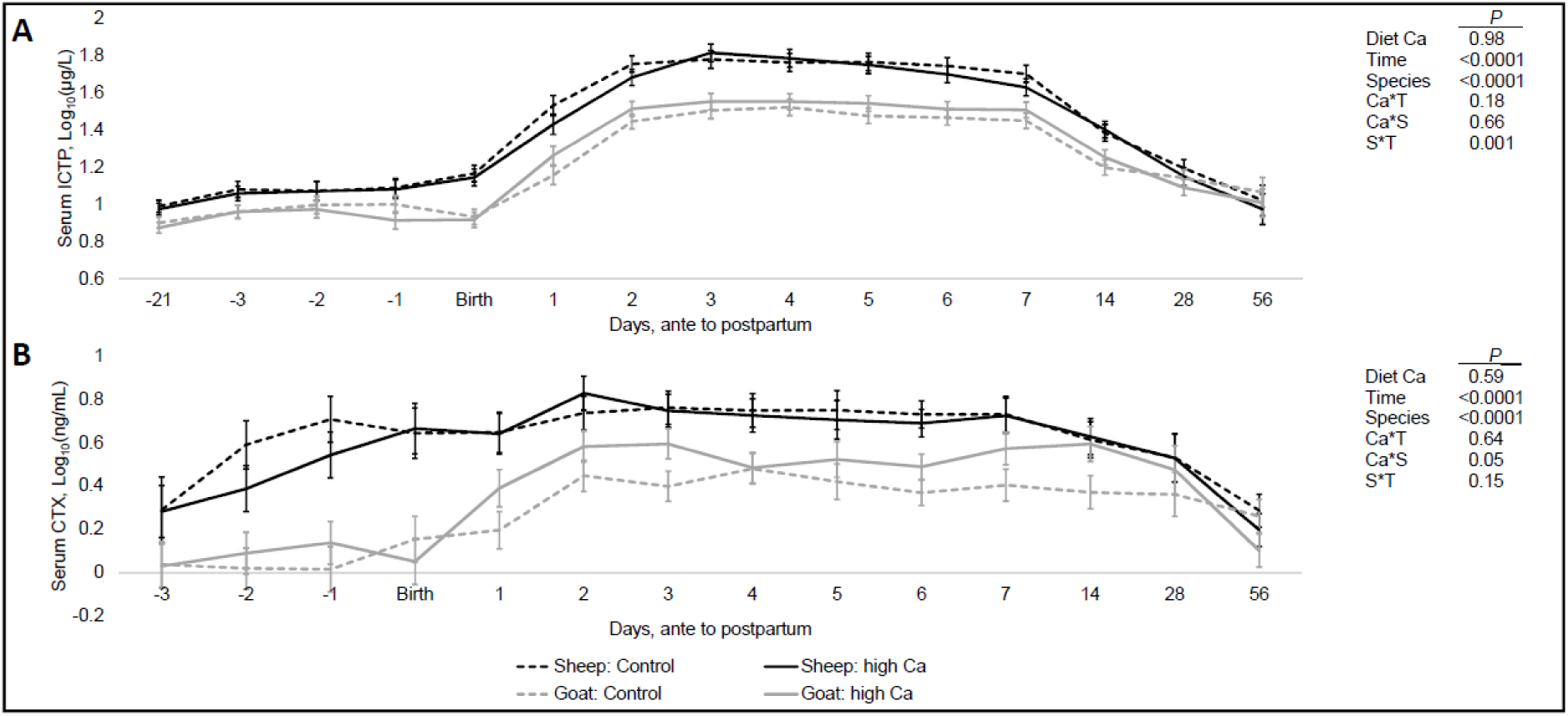
Effects of varying dietary Ca (0.6 vs. 1.3% in DM) of dairy sheep and goats during the last 3 weeks antepartum as well as time and animal species on serum ICTP (A) and serum CTX (B) from 3 weeks and 3 days antepartum, respectively, to 56 days postpartum. Effective sample size was 5 sheep and 6 goats per group. Data is presented as least square mean values (Log_10_-scale in case of bAP activity) with standard errors based on repeated measures linear mixed model analysis. Ca*T, Ca*S and S*T refer to statistical interactions between diet Ca and animal species or time, respectively, as well as animal species and time. *P* ≤ 0.05 was defined as statistical type I error threshold. Due to compromised sub-samples of analysis of CTX at −21 days antepartum and for ICTP and CTX at −4 days antepartum, we cannot provide the respective data.

Sheep showed markedly higher serum CTX levels during this study than goats (*P* < 0.0001) (Figure 6B). In sheep serum levels significantly increased from −3 days antepartum until 2 days postpartum and held a plateau until day 7 postpartum with a subsequent decline during the following weeks approaching initial values at day 57 postpartum. In contrast, goats showed not much response until parturition, after which they peaked at 2 days postpartum and fluctuated around that plateau for the next 14 days to decline until day 56 with the high Ca group showing the steepest decline. These responses over time were significant (*P* < 0.0001) but did not statistically interact with other variables like diet Ca or species. Neither on average (0.35 Cl[0.27, 0.43] vs. 0.38 Cl[0.30, 0.46] log_10_(µg/L) at 1.3% vs. 0.6% in DM; *P* = 0.59) nor over time (Diet Ca*Time: *P* = 0.64) did the feeding antepartum affect serum CTX levels in a significant manner.

### Caprine colonic abundance of mucosal vitamin D receptor

Figure 7 highlights the statistical evaluation of VDR immune reactivity in caprine colonic mucosa in goats fed 0.6% in DM and 1.3% in DM dietary Ca antepartum. Figure 8 presents exemplary immunohistochemical images from both groups at parturition. Supplementary Table 8 highlights all LSmeans and associated Tukey-adjusted 95% confidence limits of VDR immune reactivity, as related to time and dietary Ca supply antepartum.

**Figure 7.**
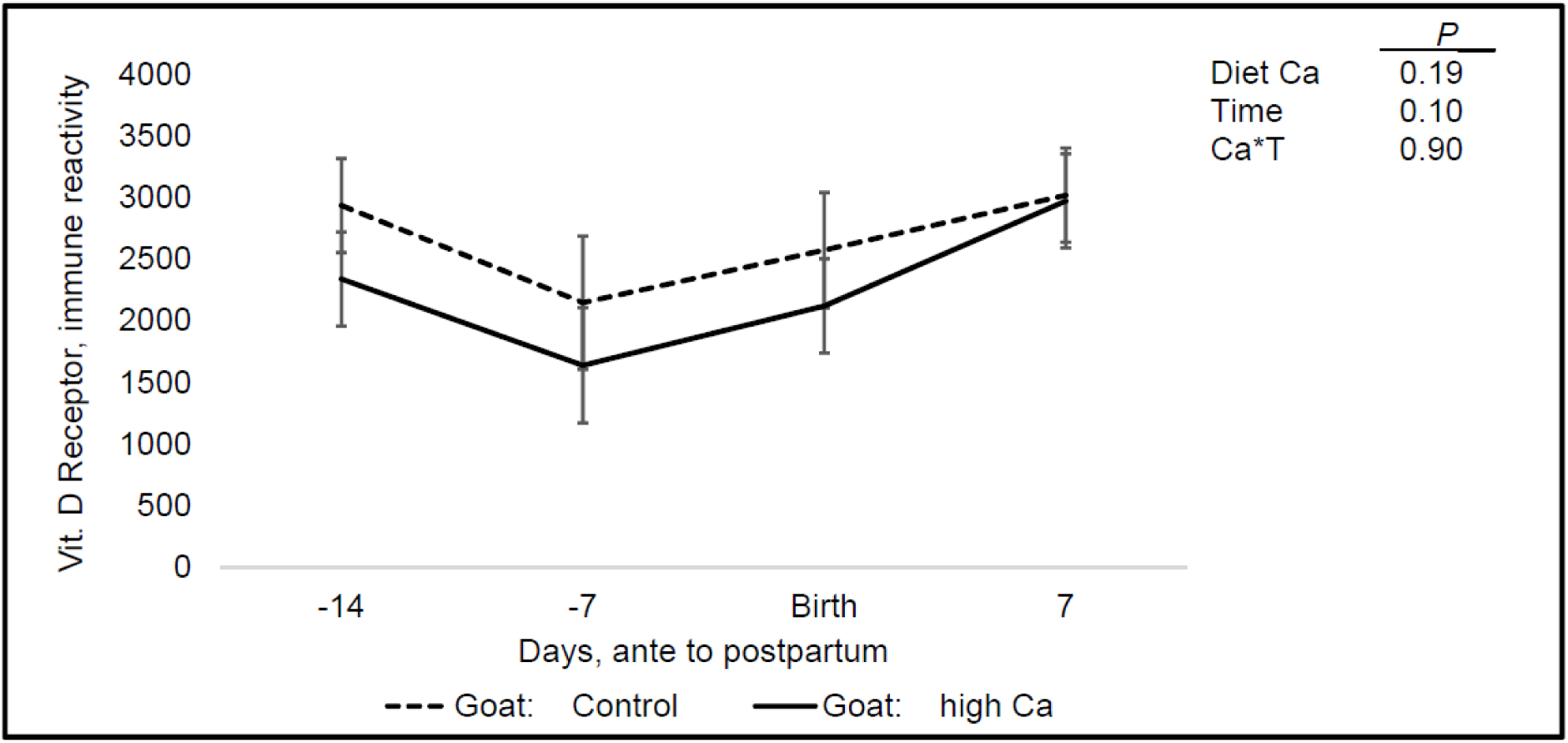
Effects of varying dietary Ca (0.6 vs. 1.3% in DM) of dairy sheep and goats during the last 3 weeks antepartum as well as time on caprine vitamin D receptor abundance in colonic mucosa from 2 weeks antepartum to 7 days postpartum. Effective sample size was 6 goats per group. Data is presented as least square mean values with standard errors based on repeated measures linear mixed model analysis. Ca*T, refers to statistical interaction between diet Ca and time. *P* ≤ 0.05 was defined as statistical type I error threshold. See Figure 8 for exemplary immunohistochemical images.

**Figure 8.**
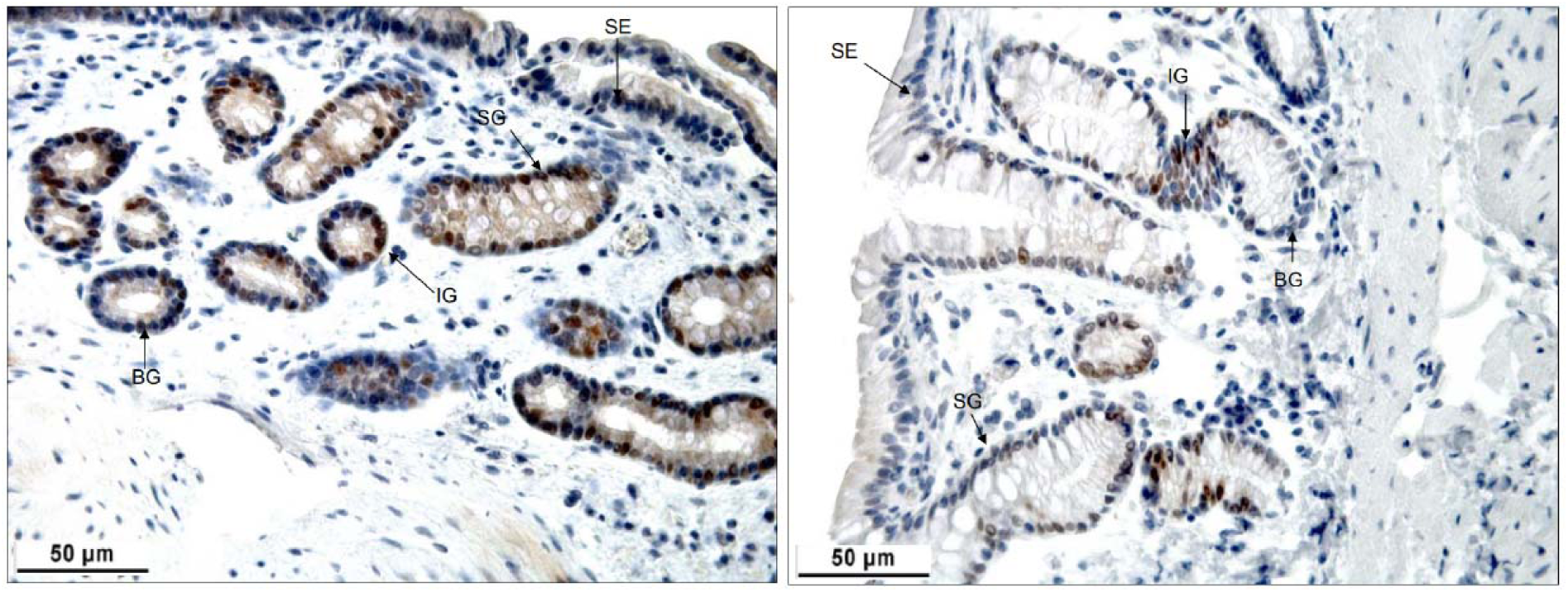
Exemplary immunohistochemical staining of the caprine VDR in the colon of goats at parturition, fed 0.6 % in DM (left) or 1.3 % in DM (right) dietary Ca during the last 3 weeks antepartum. Arrows mark SE, surface epithelium; SG, surface gland sections; IG, intermediate gland sections; BG, basal gland sections. Samples shown in these photos were taken at parturition with an endoscope equipped with biopsy forceps 40 cm into the large intestine from the anus. Goats fed high Ca antepartum showed reduced immune reactivity to VDR staining, though this could not be statistically confirmed. Figure 7 shows the statistical analysis of VDR immune reactivity over time from −14 days antepartum to 7 days postpartum.

Vitamin D receptor abundance in caprine colonic mucosa was not significantly affected by diet Ca antepartum (2268 Cl[1856, 2680] vs. 2669 Cl[2214, 3125] immune reactivity at 1.3% vs. 0.6% in DM; *P* = 0.19). In both groups, the scores showed a curvilinear trend over time which could not be statistically confirmed (*P* = 0.1).

## Discussion

An earlier meta-analysis concluded that the risk for hypocalcemia in dairy cows significantly increases if dietary Ca during the antepartum phase is between 1.1-1.5% in DM (Lean et al., 2006). This hypothesis has not yet been tested in dairy sheep and goats, therefore the present study investigated whether Ca fed at 1.3% in DM (compared to 0.6% in DM) during the last 3 weeks antepartum increases the incidence of hypocalcemia antepartum and/or postpartum by dysregulating Ca homeostasis.

### Effects on quantitative Ca homeostasis and the incidence of hypocalcemia

Sheep and goats showed in any case significant differences in the expression of observed measures of quantitative Ca homeostasis (fecal Ca, urinary Ca excretion, colostrum Ca, serum Ca, BMD). Differences in magnitude of these parameters between both species have also been observed in several earlier comparative studies. Our observation of higher values in sheep with respect to urinary Ca excretion and colostral Ca and BMD are in line with earlier studies (Liesegang et al., 2006, 2007, Liesegang, 2008). We made the same observation with serum Ca, which agrees to Kohler et al. (2013) and Liesegang (2008), though Wilkens et al. (2014) observed the opposite with goats expressing higher values than sheep. The reason for this discrepancy is unclear at the moment. Interestingly, both species exhibited a drop in serum Ca at 3 and 4 days postpartum, respectively. Similar response patterns have recently been described in dairy cows, particularly for Zn balance and serum levels (Daniel et al., 2023). It is possible that the serum Ca pattern observed in the present study mirrors a transient drop in Ca retention at the onset of lactation, comparable to the patterns observed for Zn in dairy cows. However, due to lack of data this interpretation remains speculative for the present experimental conditions and requires confirmation in future studies.

The feeding showed mostly no or just numerical effects on quantitative measures. Increasing dietary Ca feeding during the antepartum phase significantly increased fecal Ca concentration by at least 2-fold during the last 3 weeks but did not significantly affect serum and colostral Ca concentration and urinary Ca excretion in both species at any given point during the study. Most notably, no signs of hypocalcemia or any pathology were noted antepartum or postpartum, respectively, although according to Oetzel (1988), dairy sheep and goats are expected to develop milk fever symptoms comparable to cows but at 1-3 weeks postpartum. However, since 1988, advancements in milk yields and the associated calcium losses suggest that feeding regimes promoting milk fever may now be more likely to trigger the condition and potentially earlier than originally described by Oetzel. Literature on the effects of Ca oversupply in sheep and goats is scarce, particularly with respect to the antepartum phase.

However, the only study on sheep by Braithwaite (1979) agrees with our observations. That study investigated gradual differences in dietary Ca intake in adult sheep and observed increasing fecal Ca losses. Fractional Ca absorption (% of intake) was 31.8 % at the lowest Ca supply (0.22 % in DM) and dropped to a plateau around ∼9-10% with gradually increasing diet Ca to 0.6, 1.2, and 2.4% in dietary DM, respectively. In parallel, bone Ca resorption dropped inversely to the increase in Ca intake and inevitable unregulated absorption of surplus Ca was increasingly redirected into the gastrointestinal tract. Given the response of our sheep and goats in the present study, it may be concluded that for our animals and under the present study conditions, we can reject our initial hypothesis of increased milk fever risk when feeding higher Ca in the range proposed by Lean et al. (2006) for dairy cows. It remains to be tested what the outcome would be when feeding a different Ca source, for example a source with a higher water solubility like CaCl_2_ (Goff and Horst, 1993). Neither sheep nor goats showed any relevant response in urinary Ca excretion in response to the feeding regime over time, confirming once more results from earlier studies suggesting livestock ruminants do regulate Ca primarily via the gastrointestinal tract (Herm et al., 2015, Wild et al., 2021). We assume that the animals were able to dampen uptake from the gastrointestinal tract and, in parallel, effectively rerouted unavoidable unregulated Ca uptake towards excretory pathways, as described by Braithwaite (1979). Future studies must identify the precise physiological mechanisms behind this effective compensation of dietary Ca overload. See, in this context, also our discussion below on potentially confounding factors associated with our study. Furthermore, the functional basis for the differences in fecal Ca accumulation between the 2 species remains to be investigated. Initially, goats exhibited lower fecal Ca levels at −21 days antepartum, but this trend reversed until parturition. The underlying causes are currently speculative and may involve differences in dry matter digestibility or the regulation of Ca absorption at the gut barrier. Future studies should explore the effects of high Ca feeding on gastrointestinal pH, as well as the digestive and absorptive capacities for dry matter and minerals, to provide a more comprehensive understanding of this phenomenon.

### Effects on serum calcitriol and markers of bone formation and resorption

We observed a core set of serum bone formation and resorption markers, which have been earlier established in ruminants from gestation into lactation (Liesegang et al., 2000, Liesegang et al., 2007). Observed measures of bone formation and resorption showed changes over time, that have earlier been attributed to the species-dependent adaption towards parturition and the onset and further course of lactation (Liesegang et al., 2006, 2007, Liesegang, 2008, Kohler et al., 2013, Wilkens et al., 2014). However, none of the observed parameters showed a significant response to the feeding treatment during the final weeks of the antepartum and most of the postpartum period. Consequently, the BMD was not affected, suggesting that the feeding intervention with 1.3% Ca in DM did not cause any obvious dysregulation of Ca and bone homeostasis in either species as compared to the lower Ca feeding. However, serum osteocalcin showed a significant gradual increase in concentration from d7 to d56 of lactation in animals receiving the high dietary Ca antepartum. This response was true for both species, but most pronounced in the dairy goats. Serum osteocalcin is considered a marker of bone formation, with increasing levels pointing towards rising embedding of Ca in the skeleton (Liesegang et al., 2000). This observation was puzzling given the lack of response of calcitriol as well as markers of bone formation (calcitriol, bAP) and resorption (ICTP, CTX)) and especially since the effect first occurred 8 days after the suspension of the high Ca feeding. To the best of our knowledge such a response pattern has not yet been reported for serum osteocalcin, and it raises questions on its physiological purpose. The rise in serum osteocalcin was not associated with changes in BMD (or mineral concentration; data not shown) nor did other markers of bone formation show an orchestrated response that would suggest any feeding-associated changes in bone metabolism. Therefore, it may be necessary to have a look at osteocalcin from a different perspective. Biomedical research efforts in recent years suggest a hormonal role of uncarboxylated osteocalcin beyond bone metabolism, involving among others the regulation of systemic energy metabolism (Lee et al., 2007), muscle energy metabolism (Mera et al., 2016), and brain development (Obri et al., 2018). The link to the energy metabolism is particularly intriguing. A review by Wolf (2008) on the role of the skeleton in energy homeostasis suggests a connection between osteocalcin and fat deposition. Osteocalcin-deficiency as induced by an osteocalcin^-/-^ knockout increased adiposity in mice (Lee et al., 2007). It could therefore be hypothesized that an increase in circulatory osteocalcin particularly in its uncarboxylated form may have the opposite effect and that hampering the lipolysis in adipose tissue may negatively affect milk fat contents, which would explain the lower growth response of the suckling offspring of mothers fed high Ca antepartum discussed below. Indeed, this is rather speculative and more targeted research on the matter is urgently needed. In particular, the effects of high Ca antepartum on the regulation of the *Esp* gene (coding for the embryonic stem cell-specific phosphatase) in osteoblasts is of interest, given that it has been shown to be involved in regulating the carboxylation status of osteocalcin in mice (Ducy, 2011). Furthermore, maternal lipid metabolism and particularly mammary lipid metabolism should be investigated under the present experimental conditions together with a close examination of milk yield and quality over time.

### Effects on offspring development

High Ca treatment of mothers antepartum was associated with decreased average daily weight gain of suckling lambs and kids. Since the offspring showed no differences in birth weight that were associated to the feeding of the mothers antepartum, we rule out quantitative effects on placental nutrient transfer. The most plausible explanation for reduced offspring growth during the suckling phase is an impairment of milk yield and/or quality. Feed intake was not affected in the present study and all lactating animals consumed the necessary dietary dry matter according to recommendations. At the same time, we noted differences in serum osteocalcin in animals fed high Ca antepartum, pointing towards a distinct physiological conditioning. This likely has affected the quantity and/or quality of secreted milk and therefore the growth response of their offspring, but this needs yet to be confirmed. To the best of our knowledge there is yet no data available in dairy sheep and goats that allows us a more informed discussion on this matter and future studies must put a focus on the potential effects of the present experimental conditions on milk yield and quality as well as lactation physiology (see our discussion above, on the potential connection to serum osteocalcin). In addition, a further hypothesis may deserve some attention in future studies. High Ca feeding during the last 3 weeks antepartum and the associated physiological adaption of the mothers may have promoted fetal programming in the offspring that impaired their digestive capacity for Ca (and maybe other nutrients) and growth response postpartum. The possibility of mineral nutrition during gestation to induce fetal programming in ruminants either for the benefit or disadvantage of the offspring has recently been reviewed by Anas et al. (2023) for cattle, highlighting a multitude of already established interactions as well as future routes for research on the matter. In this context, an earlier study in rats tested the effects of varying dietary Ca supply during gestation including higher supply at 1.2% in DM (Li et al., 2018).

That study observed abnormal expression of hepatic and adipose genes in the offspring as well as dyslipidemia and hepatic lipid accumulation in response. Unfortunately, no data on Ca homeostasis in rat mothers and their offspring was obtained. Nevertheless, it may be argued that high supply of a mineral over several weeks may induce epigenetic changes in the mother, which could be inherited to the offspring. It may therefore be interesting to have a closer look on the epigenetic consequences of high dietary Ca as well as opposite feeding regimes during gestation and their potential to imprint Ca homeostatic regulation and development of the offspring postpartum.

### The geographical region and other potentially confounding factors

Depending on the geographic location, Ca levels in the environment are quite variable. In Switzerland, animals are often confronted with high levels of native Ca in diets, originating from the Ca-rich geological formations, which shape the chemical composition of soil in certain areas. For example an earlier survey of Schlegel et al. (2016) found a mean Ca concentration of 7 g/kg DM in 236 samples of mixed grassland from Posieux, Switzerland, and our own experience with grass hay qualities delivered to our animal facility in the Zurich area are in line with this observation. This leads to quite relevant intake levels of native Ca from feed particularly when grass-based roughage is fed, even when mineral Ca supplements are not applied. Since the organism attempts to maintain Ca homeostasis by fine-tuning absorption, losses, and mobilization of skeletal reserves according to the environmental conditions (Martín-Tereso and Martens, 2014, Wilkens and Muscher-Banse, 2020), it may be argued that sheep and goats raised predominantly on Ca-rich roughage may be better suited to deal with a high dietary Ca challenge. However, it may also suggest that during the feeding of high dietary Ca due to active supplementation of inorganic Ca to such raw materials, a metabolic imprinting as hypothesized earlier may more likely promote the observed adverse effects on daily weight gain of suckling lambs and goat kids. Therefore, the present results must be considered with caution when it comes to the general translation into dairy sheep and goat feeding systems. More data is urgently needed also from regions with less Ca in the soil.

This study is among the earliest to compare high Ca responses between sheep and goats. While the responses of many biomarkers in our study align with previous research on bone metabolism and calcium homeostasis principles within and across these species, our study is too preliminary to draw definitive conclusions regarding species-specific effects under the current treatment conditions. Given that “species” is an intrinsic characteristic, it cannot be effectively randomized within a single study limited to 1 herd per species. However, our findings represent an initial step toward further research in this area, underscoring the need for not only replication of these experimental conditions but also replication across multiple herds to strengthen the reliability of the observed phenomena.

The current Swiss feeding recommendations for sheep and goats suggest comparable dietary Ca requirements during the last 3 weeks antepartum (Agroscope, 2017). Based on these guidelines, our study applied the same diet to both species. While these recommendations likely exceed net requirements due to built-in safety margins, it remains an open question whether current genotypes in both species respond similarly to a standardized diet. To enhance future research, it would be beneficial to revisit and, if needed, revise the calcium feeding guidelines for dairy sheep and goats individually, ensuring they reflect species-specific physiological needs more accurately.

Neither sex of offspring nor litter size were balanced in the present study, therefore it may be argued whether this imbalance may have biased the results on daily weight gain during the suckling phase. However, sex was included as a variable in the statistical model and multiple births outweighed single births in both species (only 1 single birth in sheep and 2 in goats). In addition, the random baseline of each animal nested under the respective mother was added as a random factor. We are therefore confident the results truly reflect a physiological response from the varying antepartum Ca feeding of the mothers. Nevertheless, this is the 1^st^ experiment reporting these phenomena and further replication of the study ideally in other labs is needed to confirm our findings.

### Conclusions

The present study did not find any signs of hypocalcemia in sheep and goats challenged with higher Ca during the last 21 days antepartum; however, serum osteocalcin increased during the lactation in these animals. Interestingly, although no obvious negative effects of the feeding regime were observed in the mothers, the offspring of sheep and goats fed higher Ca antepartum grew slower over time compared to those in the lower Ca groups, despite having comparable birth weights. This likely originated from effects on the physiological status of the mothers, thereby affecting milk yield and quality and should be explored in future studies – with particular emphasis on the potential role of uncarboxylated osteocalcin in milk fat deposition. Furthermore, the hypothesis that high dietary Ca antepartum may adversely affect the digestive capacity of the offspring via epigenetic imprinting represents another exciting avenue for future research. Our animals were reared in a Ca-rich environment from birth; therefore, we cannot rule out that these findings are exclusive to this situation. Follow-up studies are urgently needed and should also include observations in production regions with lower soil Ca levels.

## Notes

Raw datasets can be requested via E-mail to the corresponding author.

Supplemental material for this article is available at http://dx.doi.org/10.13140/RG.2.2.32486.33601.

This study received no external funding.

We thank Dr. med. vet. Sybil Lüthi for excellent technical assistance in sample and data collection. We further thank the technical staff in the animal and laboratory facilities of the Institute of Animal Nutrition and Dietetics, Vetsuisse-Faculty, University of Zurich, Switzerland.

All procedures used in the experiment involving animals were in accordance with the Swiss guidelines for animal welfare at the time of study and were registered and approved by the Cantonal Veterinary Office (ZH 9/2006).

The authors declare that they have no known competing financial interests or personal relationships that could have appeared to influence the work reported in this paper.

## Supporting information

revised supplemental material

