## Supplementary material for "High calcium to sheep and goats antepartum Antepartum high dietary supply of calcium affects bone homeostasis and offspring growth in dairy sheep and dairy goats": revised supplemental material

D. Brugger^1^, A. Liesegang^1^

^1^Institute of Animal Nutrition and Dietetics, Vetsuisse-Faculty, University of Zurich

The supplementary material below is associated the above highlighted submission of a revised manuscript to the journal of dairy science from February 12^th^ 2025. Replication of or reference to the material should be made by citing the final published article.

Supplementary Table 1. UFA 765 supplementary concentrate for sheep and goats components and chemical composition (manufacturer information)

| **Component** | **% DM** |
| --- | --- |
| Barley flakes | 21 |
| Corn flakes | 13 |
| Wheat flakes | 6 |
| Barley | 3 |
| Wheat | 3 |
| Corn | 3 |
| Triticale | 1.5 |
| Wheat milling by-products | 14 |
| Soybean meal (40% CP) | 9 |
| Distillers grain (dark) | 7 |
| Rapeseed expeller | 2.5 |
| Rapeseed meal | 3 |
| Corn glue | 1.5 |
| Sugar beet pulp | 1 |
| Carob pellets | 2 |
| Sugar beet molasses | 4 |
| Alfalfa | 1 |
| Barley straw | 0.5 |
| Raffination fatty acids (rapeseed, sunflower) | 1 |
| Soja oil | 0.5 |
| Wheat starch | 0.5 |
| Minerals | 2 |
| Propionic acid (1k280), mg/kg | 1000 |
| Vitamin A (3a672a), IU/kg | 7500 |
| Vitamin D3 (3a671), IU/kg | 1500 |
| Vitamin E (3a700), IU/kg | 25.0 |
| Iron (3b103), mg/kg | 15.0 |
| Copper (3b405), mg/kg | 8.00 |
| Zinc (3b603), mg/kg | 35.0 |
| Manganese (3b502), mg/kg | 30.0 |
| Iodine (3b202), mg/kg | 0.40 |
| Cobalt (3b304), mg/kg | 0.10 |
| Selenium (3b801), mg/kg | 0.1 mg/kg |
| **Nutrient** | **Concentration g/kg^2^** |
| Net energy for lactation, MJ/kg | 7.00 |
| Crude ash | 55.0 |
| Crude protein | 170 |
| Crude fat | 40 |
| Calcium | 7.0 |
| Phosphorus | 4.5 |
| Magnesium | 3.0 |

^1^Communicated by the manufacturer UFA (Union des Fédérations Agricoles). For the chemical analysis of the fed batch, see Table 1 of the main manuscript.

Supplementary Table 2. Effects of varying dietary Ca (0.6 vs. 1.3% DM) of dairy sheep and goats during the last 3 weeks antepartum as well as sex and species of offspring on the birth weight (A) and daily weight gain during the suckling phase (B) – Least square mean values and associated Tukey-adjusted 95% confidence limits

| Parameter | Species | Diet Ca | Sex | Endpoint |
| --- | --- | --- | --- | --- |
| Birth weight, kg | Lamb | 0.6% in DM | male | 5.24  Cl[4.50, 6.00] |
|  |  |  | female | 4.85  Cl[4.27, 5.44] |
|  |  | 1.3% in DM | male | 5.34  Cl[4.60, 6.08] |
|  |  |  | female | 5.07  Cl[4.40, 5,75] |
|  | Kid | 0.6% in DM | male | 4.84  Cl[4.25, 5,42] |
|  |  |  | female | 4.72  [3.55, 5.89] |
|  |  | 1.3% in DM | male | 4.87  Cl[4.13, 5.61] |
|  |  |  | female | 4.32  Cl[3.74, 4.91] |
| Daily weight gain, kg/d*animal^-1^ | Lamb | 0.6% in DM | male | 0.17  Cl[0.13. 0.22] |
|  |  |  | female | 0.15  Cl[0.12, 0.19] |
|  |  | 1.3% in DM | male | 0.14  Cl[0.10, 0.19] |
|  |  |  | female | 0.13  Cl[0.09, 0.17] |
|  | Kid | 0.6% in DM | male | 0.28  Cl[0.25, 0.31] |
|  |  |  | female | 0.26  Cl[0.19, 0.32] |
|  |  | 1.3% in DM | male | 0.25  Cl[0.21, 0.29] |
|  |  |  | female | 0.21  Cl[0.18, 0.24] |

Least square mean values and associated Tukey-adjusted 95% confidence limits (Cl[upper, lower]) originate from endpoint mixed model analysis of raw data (Diet Ca, Sex, Species, age).

Supplementary Table 3. Effects of varying dietary Ca (0.6 vs. 1.3% DM) of dairy sheep and goats during the last 3 weeks antepartum as well as time and animal species on the fecal calcium concentration (A) and urinary calcium excretion (B) from 3 weeks antepartum to parturition and 14 d postpartum, respectively – Least square mean values and associated Tukey-adjusted 95% confidence limits

|  |  |  | Days antepartum | | | |  |  |  |  |  |  |  |  |  |  |
| --- | --- | --- | --- | --- | --- | --- | --- | --- | --- | --- | --- | --- | --- | --- | --- | --- |
| Parameter | Species | Diet Ca | -21 | -2 | -1 | Birth |  |  |  |  |  |  |  |  |  |  |
| Fecal Ca, % in DM | Sheep | 0.6% in DM | 2.11  Cl[1.83, 2.43] | 1.74  Cl[1.17, 2.59] | 1.75  Cl[1.20, 2.55] | 1.61  Cl[1.17, 2.21] |  |  |  |  |  |  |  |  |  |  |
|  |  | 1.3% in DM | 2.70  Cl[2.34, 3.10] | 4.10  Cl[2.93, 5,74] | 4.02  Cl[2.76, 5.86] | 4.13  [3.01, 5.67] |  |  |  |  |  |  |  |  |  |  |
|  | Goat | 0.6% in DM | 1.58  Cl[1.39, 1.80] | 3.11  Cl[2.30, 4.19] | 2.30  Cl[1.63, 3.25] | 2.24  Cl[1.68, 2.99] |  |  |  |  |  |  |  |  |  |  |
|  |  | 1.3% in DM | 1.70  Cl[1.49. 1.93] | 7.15  Cl[5.46, 9.37] | 4.63  Cl[3.16, 6.78] | 4.58  Cl[3.43, 6.11] |  |  |  |  |  |  |  |  |  |  |
|  |  |  | Days ante and postpartum | | | | | | | | | | | | | |
| Parameter | Species | Diet Ca | -21 | -4 | -3 | -2 | -1 | Birth | 1 | 2 | 3 | 4 | 5 | 6 | 7 | 14 |
| Urinary Ca excreation,  mmol/mmol creatin | Sheep | 0.6% in DM | 0.06  Cl[0.02, 0.15] | 0.03  [0.009, 0.13] | 0.04  Cl[0.01, 0.15] | 0.04  Cl[0.01, 0.14] | 0.03  Cl[0.01, 0.08] | 0.04  Cl[0.01, 0.12] | 0.05  Cl[0.02, 0.12] | 0.06  Cl[0.02, 0.16] | 0.08  Cl[0.03, 0.23] | 0.06  Cl[0.03, 0.165] | 0.05  Cl[0.02, 0.14] | 0.05  Cl[0.02, 0.13] | 0.07  Cl[0.03, 0.17] | 0.08  Cl[0.03, 0.20] |
|  |  | 1.3% in DM | 0.09  Cl[0.04, 0.24] | 0.06  Cl[0.02, 0.18] | 0.03  Cl[0.01, 0.10] | 0.04  Cl[0.02, 0.12] | 0.03  Cl[0.01, 0.07] | 0.05  Cl[0.02, 0.12] | 0.06  Cl[0.02, 0.15] | 0.04  Cl[0.02, 0.10] | 0.05  Cl[0.02, 0.12] | 0.07  Cl[0.03, 0.18] | 0.06  Cl[0.02, 0.14] | 0.04  Cl[0.02, 0.10] | 0.06  Cl[0.02, 0.15] | 0.03  Cl[0.01, 0.07] |
|  | Goat | 0.6% in DM | 0.12  Cl[0.05, 0.28] | 0.24  Cl[0.09, 0.59] | 0.34  Cl[0.15, 0.81] | 0.39  Cl[0.16, 0.98] | 0.29  Cl[0.12, 0.69] | 0.17  Cl[0.07, 0.41] | 0.09  Cl[0.04, 0.22] | 0.27  Cl[0.11, 0.64] | 0.11  Cl[0.05, 0.27] | 0.13  Cl[0.05, 0.30] | 0.12  Cl[0.05, 0.29] | 0.15  Cl[0.06, 0.36] | 0.08  Cl[0.03, 0.18] | 0.37  Cl[0.15, 0.87] |
|  |  | 1.3% in DM | 0.14  Cl[0.06, 0.33] | 0.09  Cl[0.03, 0.23] | 0.32  Cl[0.13, 0.81] | 0.13  Cl[0.06, 0.32] | 0.08  Cl[0.03, 0.20] | 0.09  Cl[0.04, 0.21] | 0.21  Cl[0.08, 0.52] | 0.11  Cl[0.04, 0.25] | 0.16  Cl[0.07, 0.39] | 0.12  Cl[0.05, 0.29] | 0.30  Cl[0.13, 0.71] | 0.17  Cl[0.07, 0.39] | 0.18  Cl[0.08, 0.43] | 0.32  Cl[0.13, 0.75] |

Least square mean values and associated Tukey-adjusted 95% confidence limits (Cl[upper, lower])originate from repeated-measures mixed model analysis of Log_10_-transformed data (Diet Ca, Time, Species).

Data was retransformed from Log_10_ to measurement scale for display in this table.

Supplementary Table 4. Effects of varying dietary Ca (0.6 vs. 1.3% DM) of dairy sheep and goats during the last 3 weeks antepartum as well as time and animal species on serum calcium (A) and colostral calcium (B) from 3 weeks antepartum to 56 d postpartum and at parturition, respectively - – Least square mean values and associated Tukey-adjusted 95% confidence limits

|  |  |  | Days ante and postpartum | | | | | | | | | | | | | | | |
| --- | --- | --- | --- | --- | --- | --- | --- | --- | --- | --- | --- | --- | --- | --- | --- | --- | --- | --- |
| Parameter | Species | Diet Ca | -21 | -4 | -3 | -2 | -1 | Birth | 1 | 2 | 3 | 4 | 5 | 6 | 7 | 14 | 28 | 56 |
| Serum Ca, mmol/L | Sheep | 0.6% in DM | 2.65  Cl[2.41, 2.89] | 2.59  Cl[2.30, 2.87] | 2.58  Cl[]2.30, 2.86 | 2.60  Cl[2.36, 2.84] | 2.61  Cl[2.28, 2.94] | 2.55  Cl[2.31, 2.79] | 2.46  Cl[2.21, 2.70] | 2.58  Cl[2.34, 2.82] | 2.74  Cl[2.50, 2.98] | 2.67  Cl[2.43, 2.92] | 2.64  Cl[2.40, 2.88] | 2.61  Cl[2.37, 2.85] | 2.68  Cl[2.44, 2.92] | 2.64  Cl[2.40, 2.88] | 2.72  Cl[2.48, 2.96] | 2.75  Cl[2.50, 2.99] |
|  |  | 1.3% in DM | 2.82  Cl[2.58, 3.07] | 2.69  Cl[2.44, 2.95] | 2.76  Cl[2.51, 3.02] | 2.70  Cl[]2.46, 2.95 | 2.80  Cl[2.48, 3.13] | 2.73  Cl[2.49, 2.97] | 2.51  Cl[2.27, 2.75] | 2.66  Cl[2.41, 2.90] | 2.37  Cl[2.12, 2.61] | 2.62  Cl[2.38, 2.86] | 2.57  Cl[2.33, 2.82] | 2.56  Cl[2.32, 2.80] | 2.60  Cl[2.36, 2.84] | 2.65  Cl[2.41, 2.89] | 2.75  Cl[2.51, 2.99] | 2.91  Cl[2.66, 3.15] |
|  | Goat | 0.6% in DM | 2.35  Cl[2.12, 2.57] | 2.36  Cl[2.13, 2.59] | 2.35  Cl[2.12, 2.59] | 2.38  Cl[2.16, 2.60] | 2.32  Cl[2.05, 2.60] | 2.26  Cl[2.04, 2.48] | 2.08  Cl[1.86, 2.30] | 1.98  Cl[1.76, 2.20] | 2.16  Cl[1.94, 2.38] | 2.24  Cl[2.01, 2.46] | 2.39  Cl[2.17, 2.61] | 2.38  Cl[2.16, 2.60] | 2.36  Cl[2.14, 2.58] | 2.29  Cl[2.07, 2.51] | 2.37  Cl[2.15, 2.59] | 2.33  Cl[2.09, 2.56] |
|  |  | 1.3% in DM | 2.42  Cl[2.19, 2.64] | 2.37  Cl[2.15, 2.59] | 2.37  Cl[2.14, 2.60] | 2.32  Cl[2.09, 2.54] | 2.43  Cl[2.20, 2.66] | 2.34  Cl[2.12, 2.56] | 2.18  Cl[1.95, 2.40] | 2.08  Cl[1.84, 2.31] | 2.07  Cl[1.85, 2.29] | 1.95  Cl[1.73, 2.17] | 2.25  Cl[2.03, 2.47] | 2.18  Cl[1.96, 2.40] | 2.22  Cl[2.00, 2.44] | 2.29  Cl[2.06, 2.51] | 2.37  Cl[2.15, 2.59] | 2.27  Cl[2.05, 2.49] |
| Parameter | Species | Diet Ca | Endpoint |  |  |  |  |  |  |  |  |  |  |  |  |  |  |  |
| Colostral Ca, g/L | Sheep | 0.6% in DM | 3.56  Cl[2.40, 4.72] |  |  |  |  |  |  |  |  |  |  |  |  |  |  |  |
|  |  | 1.3% in DM | 3.21  Cl[2.05, 4.37] |  |  |  |  |  |  |  |  |  |  |  |  |  |  |  |
|  | Goat | 0.6% in DM | 1.36  Cl[0.30, 2.42] |  |  |  |  |  |  |  |  |  |  |  |  |  |  |  |
|  |  | 1.3% in DM | 1.52  Cl[0.46, 2.57] |  |  |  |  |  |  |  |  |  |  |  |  |  |  |  |

Least square mean values and associated Tukey-adjusted 95% confidence limits (Cl[upper, lower]) originate from repeated-measures mixed model analysis (Diet Ca, Time, Species) of raw serum Ca data and endpoint mixed model analysis (Diet Ca, Species) of raw colostral Ca data.

Supplementary Table 5. Effects of varying dietary Ca (0.6 vs. 1.3% DM) of dairy sheep and goats during the last 3 weeks antepartum as well as time and animal species on bone mineral density from 2 weeks antepartum to 56 d postpartum – Least square mean values and associated Tukey-adjusted 95% confidence limits

|  |  |  | Days ante and postpartum | | | | | | |
| --- | --- | --- | --- | --- | --- | --- | --- | --- | --- |
| Parameter | Species | Diet Ca | -14 | -7 | Birth | 7 | 14 | 28 | 56 |
| Bone mineral density, mg/cm^3^ | Sheep | 0.6% in DM | 911  Cl[871, 950] | 911  Cl[838, 985] | 824  Cl[734, 915] | 859  Cl[806, 913] | 892  Cl[853, 932] | 933  Cl[903, 963] | 920  Cl[888, 953] |
|  |  | 1.3% in DM | 954  Cl[915, 994] | 903  Cl[841, 966] | 904  Cl[833, 975] | 910  Cl[860, 960] | 927  Cl[887, 967] | 962  Cl[932, 992] | 967  Cl[932, 1001] |
|  | Goat | 0.6% in DM | 860  Cl[822, 898] | 812  Cl[748, 877] | 776  Cl[715, 837] | 851  Cl[805, 896] | 875  Cl[838, 911] | 860  Cl[833, 888] | 855  Cl[826, 885] |
|  |  | 1.3% in DM | 829  Cl[788, 870] | 788  Cl[716, 859] | 788  Cl[724, 852] | 819  Cl[773, 865] | 842  Cl[806, 878] | 833  Cl[805, 860] | 836  Cl[802, 869] |

Least square mean values and associated Tukey-adjusted 95% confidence limits (Cl[upper, lower]) originate from repeated-measures mixed model analysis of raw data (Diet Ca, Time, Species).

Supplementary Table 6. Effects of varying dietary Ca (0.6 vs. 1.3% DM) of dairy sheep and goats during the last 3 weeks antepartum as well as time and animal species on serum 1.25-(OH)_2_ Vit. D, bone-specific alkaline phosphatase (bAP) activity, and osteocalcin from 3 weeks and 5 days antepartum, respectively, to 56 d postpartum – Least square mean values and associated Tukey-adjusted 95% confidence limits

|  |  |  | Days ante and postpartum | | | | | | | | | | | | | | | | |
| --- | --- | --- | --- | --- | --- | --- | --- | --- | --- | --- | --- | --- | --- | --- | --- | --- | --- | --- | --- |
| Parameter | Species | Diet Ca | -21 | | -4 | -3 | -2 | -1 | Birth | 1 | 2 | 3 | 4 | 5 | 6 | 7 | 14 | 28 | 56 |
| Serum 1.25-(OH)_2_ Vit. D., pmol/L | Sheep | 0.6% in DM | 189.6  Cl[146, 233] | | N/A | 70.4  Cl[-6.36, 147] | 113  Cl[61.6, 165] | 93.8  Cl[43.4, 144] | 76.4  Cl[50.6, 102] | 45.5  Cl[22.0, 68.9] | 65.6  Cl[8.89, 122] | 77.3  Cl[9.22, 145] | 83.9  Cl[21.1, 147] | 80.9  Cl[15.7, 146] | 80.2  Cl[8.03, 152] | 73.0  Cl[20.7, 125] | 48.0  Cl[8.54, 87.4] | 44.1  Cl[20.5, 67.7] | 69.5  Cl[36.6, 102] |
|  |  | 1.3% in DM | 133.7  Cl[90.3, 177] | | N/A | 50.4  Cl[-4.52, 105] | 59.1  Cl[13.2, 105] | 58.0  Cl[12.0, 104] | 62.6  Cl[36.8, 88.3] | 39.2  Cl[15.7, 62.7] | 72.9  Cl[16.2, 130] | 70.5  Cl[2.42, 139] | 98.5  Cl[35.8, 161] | 131  Cl[66.0, 196] | 102  Cl[29.9, 174] | 94.1  Cl[41.8, 146] | 79.6  Cl[40.2, 119] | 52.6  Cl[28.9, 76.2] | 46.4  Cl[13.5, 79.3] |
|  | Goat | 0.6% in DM | 81.7  Cl[42.1, 121] | | N/A | 100  Cl[50.9, 149] | 144  Cl[105, 184] | 145  Cl[93.5, 196] | 165  Cl[142, 189] | 154  Cl[132, 175] | 181  Cl[129, 232] | 212  Cl[150, 274] | 206  Cl[149, 263] | 164  Cl[105, 224] | 155  Cl[88.9, 221] | 140  Cl[92.4, 188] | 117  Cl[81.1, 153] | 120  Cl[98.1, 141] | 91.8  Cl[58.5, 125] |
|  |  | 1.3% in DM | 98.2  Cl[58.6, 138] | | N/A | 110  Cl[65.0, 155] | 111  Cl[70.1, 153] | 158  Cl[114, 202] | 155  Cl[132, 179] | 137  Cl[116, 159] | 196  Cl[144, 248] | 205  Cl[143, 267] | 232  Cl[174, 289] | 209  Cl[150, 269] | 218  Cl[152, 284] | 191  Cl[144, 239] | 148  Cl[112, 184] | 119  Cl[97.9, 141] | 120  Cl[89.8, 150] |
|  |  |  | Days ante and postpartum | | | | | | | | | | | | | | | | |
| Parameter | Species | Diet Ca | -21 |  | -4 | -3 | -2 | -1 | Birth | 1 | 2 | 3 | 4 | 5 | 6 | 7 | 14 | 28 | 56 |
| Serum bAP, U/L | Sheep | 0.6% in DM | 93.4  Cl[55.7, 157] | | 89.2  Cl[56.7, 140] | 95.0  Cl[57.6, 157] | 87.3  Cl[55.3, 138] | 84.7  Cl[34.4, 208] | 69.9  Cl[29.3, 166] | 72.4  Cl[49.3, 106] | 82.1  Cl[54.6, 123] | 84.6  Cl[60.1, 119] | 77.4  Cl[52.8, 113] | 75.9  Cl[50.1, 115] | 74.2  Cl[47.8, 115] | 74.3  Cl[46.4, 119] | 70.9  Cl[44.0, 114] | 76.8  Cl[45.8, 129] | 98.3  Cl[60.5, 160] |
|  |  | 1.3% in DM | 129  Cl[76.7, 215] | | 117  Cl[73.8, 186] | 99.1  Cl[59.3, 165] | 111  Cl[69.7, 177] | 106  Cl[43.2, 261] | 86.1  Cl[36.1, 205] | 104  Cl[70.8, 153] | 112  Cl[74.5, 168] | 107  Cl[76.2, 151] | 109  Cl[74.3, 160] | 104  Cl[68.9, 158] | 104  Cl[67.1, 162] | 95.6  Cl[59.7, 153] | 87.2  Cl[54.1, 141] | 96.4  Cl[57.4, 162] | 154  Cl[87.7, 271] |
|  | Goat | 0.6% in DM | 51.4  Cl[32.1, 82.3] | | 42.8  Cl[28.3, 64.7] | 41.4  Cl[26.2, 65.3] | 47.5  Cl[31.5, 71.5] | 20.6  Cl[8.85, 48.1] | 20.1  Cl[9.08, 44.4] | 31.0  Cl[21.8, 44.1] | 34.7  Cl[23.9, 50.4] | 41.7  Cl[30.5, 57.0] | 45.9  Cl[32.3, 65.1] | 46.6  Cl[31.9, 68.1] | 47.1  Cl[31.5, 70.5] | 43.8  Cl[28.5, 67.4] | 40.7  Cl[26.3, 62.9] | 43.5  Cl[27.1, 69.7] | 44.0  Cl[27.7, 70.1] |
|  |  | 1.3% in DM | 48.0  Cl[26.9, 85.4] | | 39.3  Cl[23.8, 65.1] | 37.6  Cl[21.5, 65.8] | 42.4  Cl[25.6, 70.0] | 36.0  Cl[13.0, 100] | 34.5  Cl[13.1, 91.2] | 37.4  Cl[24.4, 57.6] | 33.2  Cl[21.1, 52.3] | 30.3  Cl[20.7, 44.4] | 34.2  Cl[22.3, 52.5] | 31.8  Cl[20.0, 50.6] | 32.6  Cl[19.9, 53.3] | 41.7  Cl[24.6, 70.6] | 31.9  Cl[18.7, 54.5] | 34.1  Cl[19.1, 60.8] | 39.9  Cl[23.2, 68.7] |
|  |  |  | Days ante and postpartum | | | | | | | | | | | | | | | | |
| Parameter | Species | Diet Ca | -21 |  | -4 | -3 | -2 | -1 | Birth | 1 | 2 | 3 | 4 | 5 | 6 | 7 | 14 | 28 | 56 |
| Serum Osteocalcin, mmol/L | Sheep | 0.6% in DM | 16.4  Cl[5.53, 27.3] | | 18.4  Cl[10.7, 26.2] | 20.5  Cl[15.2, 25.8] | 19.1  Cl[11.7, 26.5] | 19.0  Cl[12.0, 26.1] | 13.8  Cl[9.56, 18.0] | 18.3  Cl[11.8, 24.7] | 15.7  Cl[10.5, 20.9] | 16.5  Cl[12.0, 20.9] | 15.7  Cl[12.0, 19.4] | 16.4  Cl[12.4, 20.4] | 13.3  Cl[4.63, 22.0] | 15.5  Cl[6.37, 24.7] | 16.2  Cl[0.99, 31.4] | 15.5  Cl[-5.63, 36.7] | 29.9  Cl[5.62, 54.1] |
|  |  | 1.3% in DM | 18.1  Cl[10.2, 26.1] | | 14.3  Cl[8.44, 20.2] | 16.5  Cl[11.7, 21.3] | 16.1  Cl[8.65, 23.5] | 22.7  Cl[15.7, 29.8] | 15.1  Cl[10.9, 19.3] | 14.5  Cl[8.03, 20.9] | 14.4  Cl[9.19, 19.6] | 15.9  Cl[11.4, 20.4] | 16.2  Cl[12.5, 20.0] | 14.2  Cl[10.2, 18.2] | 14.9  Cl[6.20, 23.6] | 15.1  Cl[5.95, 24.2] | 19.9  Cl[4.72, 35.2] | 27.8  Cl[6.61, 49.0] | 39.6  Cl[15.3, 63.8] |
|  | Goat | 0.6% in DM | 20.2  Cl[13.4, 27.0] | | 18.4  Cl[13.4, 23.4] | 18.1  Cl[13.9, 22.3] | 16.9  Cl[10.2, 23.7] | 13.4  Cl[7.30, 19.6] | 14.0  Cl[10.1, 17.8] | 14.8  Cl[8.90, 20.7] | 15.6  Cl[10.8, 20.3] | 13.2  Cl[9.11, 17.2] | 17.2  Cl[13.8, 20.6] | 17.9  Cl[14.3, 21.6] | 22.4  Cl[14.5, 30.4] | 19.2  Cl[10.8, 27.5] | 23.7  Cl[9.84, 37.6] | 23.6  Cl[4.26, 42.9] | 42.2  Cl[17.8, 66.6] |
|  |  | 1.3% in DM | 16.2  Cl[8.90, 23.4] | | 18.6  Cl[13.8, 23.4] | 15.9  Cl[11.7, 20.1] | 20.5  Cl[13.7, 27.3] | 16.1  Cl[10.3, 21.9] | 14.4  Cl[10.6, 18.3] | 19.4  Cl[13.5, 25.3] | 15.8  Cl[11.0, 20.5] | 18.2  Cl[14.1, 22.3] | 17.0  Cl[13.6, 20.4] | 19.6  Cl[16.0, 23.3] | 26.7  Cl[18.8, 34.7] | 33.4  Cl[25.1, 41.7] | 39.2  Cl[25.3, 53.1] | 73.8  Cl[53.9, 93.6] | 80.6  Cl[58.4, 103] |

Least square mean values and associated Tukey-adjusted 95% confidence limits (Cl[upper, lower]) originate from repeated-measures mixed model analysis (Diet Ca, Time, Species) of raw serum 1.25-(OH)_2_ Vit. D and osteocalcin data as well as Log_10_-transformed bAP data.

bAP Data was retransformed from Log_10_ to measurement scale for display in this table.

N/A, not assessed due to compromised sample quality.

Supplementary Table 7. Effects of varying dietary Ca (0.6 vs. 1.3% DM) of dairy sheep and goats during the last 3 weeks antepartum as well as time and animal species on serum crosslinked carboxaterminal teleopeptide of type I collagen (ICTP) and crosslaps (CTX) from 3 weeks and 3 days antepartum, respectively, to 56 d postpartum – Least square mean values and associated Tukey-adjusted 95% confidence limits

|  |  |  | Days ante and postpartum | | | | | | | | | | | | | | |
| --- | --- | --- | --- | --- | --- | --- | --- | --- | --- | --- | --- | --- | --- | --- | --- | --- | --- |
| Parameter | Species | Diet Ca | -21 | 3 | -2 | -1 | Birth | 1 | 2 | 3 | 4 | 5 | 6 | 7 | 14 | 28 | 56 |
| Serum ICTP, µg/L | Sheep | 0.6% in DM | 9.77  Cl[8.38, 11.4] | 12.0  Cl[9.79, 14.8] | 11.8  Cl[9.21, 15.2] | 12.3  Cl[9.60, 15.7] | 14.7  Cl[11.8, 18.2] | 34.0  Cl[26.1, 44.3] | 56.9  Cl[46.1, 70.2] | 60.2  Cl[47.7, 76.1] | 58.0  Cl[46.1, 73.0] | 58.4  Cl[46.6, 73.1] | 55.6  Cl[44.8, 68.9] | 50.4  Cl[40.4, 62.9] | 24.2  Cl[19.5, 30.1] | 15.7  Cl[12.5, 19.7] | 10.5  Cl[7.06, 15.6] |
|  |  | 1.3% in DM | 9.44  Cl[8.10, 11.0] | 11.4  Cl[9.46, 13.8] | 11.8  Cl[9.19, 15.1] | 12.1  Cl[9.44, 15.4] | 14.0  Cl[11.3, 17.3] | 26.9  Cl[20.7, 35.1] | 48.1  Cl[39.0, 59.4] | 65.3  Cl[51.7, 82.5] | 61.1  Cl[48.5, 76.9] | 56.3  Cl[44.9, 70.4] | 50.0  Cl[40.3, 62.0] | 42.6  Cl[34.1, 53.1] | 25.2  Cl[20.3, 31.3] | 14.1  Cl[11.3, 17.7] | 9.41  Cl[6.33, 14.0] |
|  | Goat | 0.6% in DM | 7.97  Cl[6.93, 9.17] | 9.13  Cl[7.69, 10.8] | 9.94  Cl[7.92, 12.5] | 9.99  Cl[7.97, 12.5] | 8.58  Cl[7.05, 10.4] | 14.3  Cl[11.2, 18.2] | 27.9  Cl[23.0, 33.8] | 32.0  Cl[25.9, 39.7] | 33.3  Cl[27.0, 41.0] | 30.0  Cl[24.4, 36.8] | 29.3  Cl[24.0, 35.6] | 28.1  Cl[23.0, 34.5] | 15.8  Cl[13.0, 19.2] | 13.9  Cl[11.3, 17.0] | 11.6  Cl[7.91, 17.1] |
|  |  | 1.3% in DM | 7.50  Cl[6.51, 8.62] | 9.14  Cl[7.71, 10.8] | 9.41  Cl[7.50, 11.8] | 8.20  Cl[6.56, 10.3] | 8.27  Cl[6.79, 10.1] | 18.3  Cl[14.4, 23.3] | 32.6  Cl[26.9, 39.5] | 35.8  Cl[28.9, 44.2] | 35.7  Cl[28.9, 44.0] | 34.9  Cl[28.4, 42.8] | 32.5  Cl[26.7, 39.6] | 32.2  Cl[26.3, 39.4] | 17.9  Cl[14.7, 21.8] | 12.3  Cl[10.0, 15.2] | 10.2  Cl[7.09, 14.6] |
|  |  |  | Days ante and postpartum | | | | | | | | | | | | | | |
| Parameter | Species | Diet Ca | -21 | 3 | -2 | -1 | Birth | 1 | 2 | 3 | 4 | 5 | 6 | 7 | 14 | 28 | 56 |
| Serum CTX, ng/mL | Sheep | 0.6% in DM | N/A | 1.95  Cl[0.94, 4.02] | 3.89  Cl[2.29, 6.63] | 5.10  Cl[3.05, 8.51] | 4.40  Cl[2.51, 7.72] | 4.44  Cl[2.82, 6.98] | 5.45  Cl[3.74, 7.93] | 5.77  Cl[3.99, 8.35] | 5.60  Cl[3.86, 8.13] | 5.62  Cl[3.64, 8.66] | 5.37  Cl[3.96, 7.30] | 5.37  Cl[3.62, 7.97] | 4.11  Cl[2.74, 6.16] | 3.39  Cl[1.98, 5.81] | 1.94  Cl[1.33, 2.81] |
|  |  | 1.3% in DM | N/A | 1.92  Cl[1.07, 3.43] | 2.44  Cl[1.46, 4.08] | 3.49  Cl[2.09, 5.83] | 4.62  Cl[2.63, 8.10] | 4.37  Cl[2.78, 6.88] | 6.73  Cl[4.62, 9.79] | 5.58  Cl[3.85, 8.07] | 5.31  Cl[3.66, 7.71] | 5.08  Cl[3.29, 7.83] | 4.89  Cl[3.60, 6.65] | 5.30  Cl[3.57, 7.87] | 4.25  Cl[2.83, 6.37] | 3.37  Cl[1.97, 5.78] | 1.57  Cl[1.09, 2.28] |
|  | Goat | 0.6% in DM | N/A | 1.09  Cl[0.66, 1.82] | 1.05  Cl[0.66, 1.65] | 1.04  Cl[0.63, 1.71] | 1.43  Cl[0.86, 2.39] | 1.57  Cl[1.04, 2.38] | 2.80  Cl[1.99, 3.94] | 2.50  Cl[1.79, 3.51] | 3.02  Cl[2.15, 4.24] | 2.63  Cl[1.77, 3.90] | 2.34  Cl[1.77, 3.09] | 2.53  Cl[1.77, 3.63] | 2.35  Cl[1.62, 3.40] | 2.30  Cl[1.41, 3.75] | 1.82  Cl[1.26, 2.65] |
|  |  | 1.3% in DM | N/A | 1.07  Cl[0.65, 1.77] | 1.23  Cl[0.77, 1.96] | 1.37  Cl[0.85, 2.21] | 1.13  Cl[0.68, 1.88] | 2.46  Cl[1.62, 3.71] | 3.82  Cl[2.71, 5.39] | 3.93  Cl[2.80, 5.51] | 3.05  Cl[2.17, 4.29] | 3.33  Cl[2.24, 4.94] | 3.08  Cl[2.33, 4.07] | 3.73  Cl[2.60, 5.35] | 3.92  Cl[2.66, 5.78] | 2.98  Cl[1.74, 5.11] | 1.27  Cl[0.87, 1.84] |

Least square mean values and associated Tukey-adjusted 95% confidence limits (Cl[upper, lower]) originate from repeated-measures mixed model analysis of Log_10_-transformed data (Diet Ca, Time, Species).

Data was retransformed from Log_10_ to measurement scale for display in this table.

N/A, not assessed due to compromised sample quality.

Supplementary Table 8. Effects of varying dietary Ca (0.6 vs. 1.3% DM) of dairy sheep and goats during the last 3 weeks antepartum as well as time and animal species on vitamin D receptor immune reactivity from 2 weeks antepartum to 7 d postpartum – Least square mean values and associated Tukey-adjusted 95% confidence limits

|  |  |  | Days ante and postpartum | | | |
| --- | --- | --- | --- | --- | --- | --- |
| Parameter | Species | Diet Ca | -14 | -7 | Birth | 7 |
| Vitamin D receptor, immune reactivity | Goat | 0.6% in DM | 2936  Cl[2159, 3713] | 2148  Cl[1049, 3247] | 2574  Cl[1622, 3526] | 3019  Cl[2242, 3796] |
|  |  | 1.3% in DM | 2340  Cl[1562, 3117] | 1639  Cl[687, 2591] | 2121  Cl[1344, 2899] | 2973  Cl[2196, 3750] |

Least square mean values and associated Tukey-adjusted 95% confidence limits (Cl[upper, lower]) originate from repeated-measures mixed model analysis of raw data (Diet Ca, Time).
